# MERIT: Mechanism driven model predicts drug outcomes and nominates indications for failed drugs

**DOI:** 10.64898/2026.08.09.743088

**Authors:** H.H. Caline Koh-Tan, Ivan Meić, Buğra Alptekin Sarı, Steve Muller, Gabriel Richman

## Abstract

Drug development depends on efficacy and safety, but many trial-outcome prediction models incorporate trial design, prior development history or compound identity, enabling compound memorization and inflating apparent performance. We developed MEchanism-Resolved Inference of Trial outcomes (MERIT), a model that predicts trial outcomes from molecular and disease features without using information on similar-compound success. MERIT integrates the disease and drug of interest with large-scale drug-protein, protein-metabolite and immune interaction maps to link a drug’s intended and potential off-target effects to tissue-specific efficacy and safety. Across 753 small-molecule drugs and 3,133 trials, MERIT achieved an overall AUROC of 0.770 (0.765 for efficacy and 0.784 for safety) under structure-holdout evaluation, outperforming two published methods retrained on the same trials and folds. Scored by a mechanism-fit profiling, MERIT ranked the approved indication above the failed one for 83% (50 of 60) of a preselected set of 60 cohort compounds that failed one indication but are approved for another. Finally, we registered locked, outcome-blind predictions for 55 drug-indication pairs in ongoing phase III trials for prospective evaluation.

## INTRODUCTION

When a drug fails in the clinic, the cause may be a property of the molecule or external to it (such as a trial stopped over funding, enrollment or corporate priorities)^1^. Only the former reflects a property of the compound, yet the two are seldom separated. Among genuine biological failures, safety failures arise from off-target pharmacology, metabolic activation and organ-specific toxicity^2^, whereas efficacy failures, predominant cause of late-stage drug failure, reflect a mechanism that does not engage disease biology^3^. Whether these drug-driven outcomes can be predicted from a compound’s biology before development is committed has remained open^4^.

Computational models have predicted trial-related outcomes for years, reporting area-under-receiver-operator-curve (AUROCs) of roughly 0.72 to 0.88^5–8^. Most of these numbers come from evaluations in which the very same drug appears in both training and test datasets^9^. A model can score well from the recorded outcomes of specific drugs, without information on why a drug’s biology suits a disease. A model that has already seen aspirin succeed can predict its next trial outcome from aspirin’s identity alone. A model is only informative for a development decision if it can predict a novel drug’s outcome.

To objectively evaluate model performance, predictions should be compared under two evaluation conditions. In the strict evaluation, every drug in the test set is chemically new to the model. In the loose evaluation, the drug’s other trials are allowed to remain in training. The difference between the two defines the memorization gap, the share of apparent accuracy that comes from the presence of the same drug’s other trials in training rather than from its biology.

This study introduces the MEchanism-Resolved Inference of Trial outcomes (MERIT), a model that predicts trial outcomes and ranks candidate indications from a drug’s biology rather than its identity. This study profiled 753 small-molecule drugs across their 3,133 trials records, combining disease context with molecular-mechanism profiling. Previously reported models scored lower under a strict structure-holdout test that excludes drugs found in the training dataset. Retrained on the same trials, MERIT achieved an AUROC of 0.078 above a leading model HINT^6^, and 0.131 above TrialBench^7^. MERIT achieved a structure-holdout AUROC of 0.770, contributed primarily by two pre-trial factors, disease complexity and match between drug’s mechanism of action and the target indication.

## METHODS

### Operational definitions

**PASS / “success”** – Completion of a phase III/IV trial with no recorded termination, as reported on ClinicalTrials.gov. This is not primary-endpoint success, approval, or demonstrated efficacy. Phase I/II completions are labeled IN_PROCESS and excluded.

**FAIL_EFFICACY / “efficacy failure”** – A terminated trial where blind audit attributes the stop to lack of efficacy or futility of the investigational compound. This is not a finding that the drug is inefficacious in general.

**FAIL_SAFETY / “safety failure”** – A terminated trial where blind audit attributes the stop to a drug-related toxicity or safety finding of the investigational compound, excluding toxicity attributable to another combination drug.

**FAIL_BOTH** – Both efficacy and safety failures present (n = 5).

**“Failure sets”** – Any trial that failed efficacy, safety or both.

**“Structure-holdout” / “Strict” evaluation** – Cross-validation grouped by InChIKey connectivity layer (IK14) so no compound appears in both training and test datasets.

**“Structure-blind”** / **“Loose”** evaluation assigns the same trials to folds at random.

**“Memorization gap”** – The increase in AUROC when a compound’s other trials are in training dataset. The compound-level features have the same values in every trial of a compound, so the model can recognize the compound and reuse the outcomes of its other trials, instead of predicting from its biology. The gap is the AUROC under structure-blind evaluation minus the AUROC under structure-holdout evaluation.

**“Mechanism-fit”** – The quantity [1 − P(efficacy failure)] from the model efficacy classifier computed with the compound held out of training. A model-derived score of how well the compound’s target biology matches the indication. This is not a measure of clinical effect size or a probability of approval.

**“Novel drug-indication pair”** – A cohort compound paired with an indication and this pairing was not tested previously, anywhere in the training cohort.

**“Prospective”** – Used in two contexts, distinguished at each use: (i) *temporal (leave-future-out) validation*, with retrospective trials split by start date, where outcomes were already known at analysis time; and (ii) *prospective registration*, the 55 predictions locked for ongoing phase III trials with outcomes unknown to us at the lock date.

### Data sources and label verification

Clinical trial data were obtained from the Aggregate Analysis of Clinical Trials (AACT) database^10^ on 2-Apr-2026, comprising 428,377 drug-intervention records. After filtering to drug-type interventions with clear outcomes, 12,420 trials across 2,048 unique drug names were identified. SMILES structures were resolved for 1,886 drugs through PubChem^11^ (primary), ChEMBL^12^ (secondary) and manual curation, while 162 drugs were unresolvable (biologics, experimental codes, non-drug entries) and explicitly categorized. SMILES were standardized via RDKit^13^, salt-stripped, neutralized, and deduplicated by the International Chemical Identifier key (InChIKey) connectivity layer (first 14 characters) to IK14, collapsing salt forms while preserving stereoisomers as distinct compounds.

All 672 terminated trials were classified from the *why_stopped* field using keyword matching with priority ordering (safety > efficacy > enrollment > business), and subsequently corrected by manual review. This corrected 34 misclassifications, removed 15 non-drug entries (antibody-drug conjugates, peptides, combination regimens, plant supplements) and corrected 4 misidentified compounds.

Disease labels were manually curated where the original AACT conditions were missing or ambiguous. Procedural/anesthetic trials were excluded from efficacy evaluation but retained for safety. Anti-pathogen drugs were excluded from efficacy evaluation because the binding profile predicts human proteome interactions, not drug-pathogen interactions (***Table S1***). A blind, web-verified label audit reclassified and removed non-drug stops from the any of the FAIL groups. The trial-level modeling cohort comprises 3,133 trial records across 753 compounds (2,967 distinct trials, where a distinct trial tested several compounds contributing to one record per compound) with a complete molecular-mechanism profile (PASS 2,558; FAIL_EFFICACY 490; FAIL_SAFETY 80; FAIL_BOTH 5). The arm-level robustness cohort (***Table S2***) comprises 3,326 arms across 891 compounds. Compounds lacking a complete computational profile (predominantly large molecules) were excluded.

### Computational modules

The candidate feature pool comprises 285 features across nine modules. The tissue interaction module (module 1) contributes 97 features that weigh predicted binding by tissue-specific expression from the Human Protein Atlas^14^. The network-pathway enrichment module (module 2) contributes 27 features that map binding targets onto STRING^15^ protein-protein interaction networks and quantify enrichment against Open Targets^16^ disease genes. Binding specificity (module 3) contributes 30 features. Safety pharmacology (module 4) adds 45 features. A further 10 pharmacokinetic features (module 5) are predicted with DruMAP^17^. The target-disease biology module (module 6), contributes 30 features computed from Open Targets biology channels, ChEMBL^12^ mechanism-of-action annotation, ClinGen^18^ germline genetics, STRING network and KEGG^19^ pathway membership. The genetics module (module 7) contributes 16 features from and DepMap^20^ dependency and ClinVar^21^ germline genetics. Disease complexity and trial context (module 8) accounts for 15 features. The trial-design module (module 9) adds the remaining 15 features. All module feature enumerations are given in ***Notes S1 and S2***, and the released *feature_list_final.csv* gives the exact module-family assignment for every candidate feature.

For each cross-validation training fold, the top 20 candidate features with the highest individual AUROC against the outcome are selected and used to fit that fold’s classifier.

### Model training and evaluation

MERIT’s classifiers are gradient-boosted decision trees with 500 trees, with maximum depth of 3, a learning rate of 0.05 and a subsample fraction of 0.8. The overall task and each safety detector use a single scikit-learn gradient-boosting algorithm, whereas the efficacy task uses an equal-weight average of three gradient-boosting implementations (scikit-learn^22^, XGBoost^23^ and LightGBM^24^) fitted with balanced class weights (random seeds 42, 123, 456, 789, 2024). StratifiedGroupKFold cross-validation with SMILES as the grouping variable ensures zero compound overlap between train and test datasets, repeated across 5 random seeds × 5 folds for 25 evaluations. Feature selection and median imputation (fitted on training data only) prevent information leakage. The primary analysis is at the trial level, repeated at the individual investigational arm under the same structure-holdout protocol (***Table S2***). All AUROC values are reported as the mean ± SD across the 25 folds. These 25 evaluations are not fully independent since the 5 seeds reshuffle the same trials so the SD understates between-sample variability. Events-per-variable on the 20 selected features is 29 overall, 25 for efficacy and 4 for safety, so safety is power-limited. The analysis uses scikit-learn 1.6.1, XGBoost 3.0.3 and LightGBM 4.6.0 under Python 3.12.

#### Classification metrics

AUROC, accuracy, balanced accuracy, precision, recall, F1, Matthews correlation coefficient (MCC) and precision-recall AUC (PR-AUC) are computed per fold and averaged over the 25 evaluations, at two operating points of default 0.50 threshold and each task’s training-fold base rate. The base rate is the operating point relevant to prioritizing candidates given class imbalance. The overall and efficacy tasks use calibrated probabilities, whereas safety uses an uncalibrated ranker (***Table S4***).

#### Probability calibration

Per-fold isotonic regression was applied to raw out-of-fold (OOF) probabilities using a 3-fold nested split within each outer training fold (no label leakage across the outer structure-holdout boundary). Isotonic calibration reduced the pooled-OOF expected calibration error (ECE) from 0.097 to 0.035 (overall) and 0.078 to 0.032 (efficacy), with the fold-averaged AUROC unchanged. The two tasks are binned identically, on seed-averaged out-of-fold calibrated probabilities over the task’s own cohort (***Table S5***).

#### Traditional machine-learning baselines

Random forest (500 trees, minimum leaf size 2, balanced class weights), logistic regression (L2, balanced class weights, standardized inputs) and a single gradient-boosting algorithm were each trained and evaluated separately on the overall, efficacy and safety tasks. All three algorithms used the same feature set, under identical cohorts, labels, folds, median imputation, the same per-fold top-20 feature selection and within-fold disease encoding, with only the algorithm substituted, and scored on the same metric panel (***Table S6***). The comparison for the efficacy task uses the single gradient-boosting algorithm instead of MERIT’s three gradient-boosting implementations. Label permutation testing (50 shuffles) used the same compound-based fold assignment to test whether apparent signal arises from chance fold assignment.

#### Benchmark comparisons

MERIT was compared with two published methods, HINT^8^ and and TrialBench^7^. HINT is a deep-learning model (molecular graph, disease codes and eligibility-criteria text), and TrialBench is a multi-modal benchmark of trial-design features. The TrialBench comparison used the trial-identifier intersection of this study’s cohort with TrialBench’s trial-approval task (1,142 trials, 412 compounds). The TrialBench approval label was predicted under the same structure-holdout (IK14 grouping; 3 seeds × 5 folds; gradient-boosting). This protocol score TrialBench’s tabular trial-design features, MERIT’s mechanism-based features and the union. The same feature restrictions apply to both methods. MERIT’s feature set excludes trial-design and trial-context features, and the TrialBench set is reduced to trial-design features only by removing the enrollment count and the establishment metadata. The published HINT was first reproduced on its own published benchmark to confirm it matches the released HINT predictions (***Notes S3***). It was then retrained on this study’s cohort using the same structure-holdout folds as MERIT (2,787 unique-NCT, IK14-grouped folds).

#### Network-medicine baseline

The closest-distance network proximity between each drug’s protein targets and the disease gene module was computed over the STRING v12 interactome (combined-score ≥ 400; 19,488 nodes)^25^. Each proximity (d_c) was z-scored against 50 degree-preserving random gene-set pairs, then scored directly as an efficacy predictor on the trials with both annotations (1,385 trials, 223 failures) and evaluated by drug-clustered AUROC.

#### The memorization gap (evaluation-inflation analysis)

Each feature set was scored under two cross-validation evaluations differing only in fold assignment. Cohort, label, algorithm, imputation, feature selection, seeds and fold count are constant between the two evaluations. The structure-holdout evaluation groups folds by the IK14 (StratifiedGroupKFold), so no compound appears in both training and test dataset, whereas the structure-blind evaluations assigns the same trials to folds at random. The inflation is the difference in fold-averaged AUROC between the two evaluations. The structure-holdout evaluation removes the test compound from the downstream classifier, but the upstream structure-to-target binding predictor is separately trained, so a compound with abundant public bioactivity data could have a more informative binding profile. Three analyses address this limitation. First, the binding profiles come from a pretrained drug-target affinity model without trial outcomes or clinical annotations. Second, a public-only feature set was evaluated under identical protocol to isolate how much of the efficacy increment remains without it. Third, the binding-derived safety classifier was also evaluated for correlation with black-box-warnings or post-market-withdrawal statuses on the reasoning that a classifier using public information on how established a drug is instead of structure would be associated with those labels.

#### Safety classifier

The safety classifier is a single gradient-boosted algorithm over the union of the six safety axes (promiscuity, hepatic, cardiac, network, tissue, context), trained under the identical structure-holdout folds. It was compared against a parameter-free noisy-OR, the probabilistic form of a logical disjunction (OR) over six independent per-axis detectors [P(fail) = 1 − Π_m (1 − p_m)]. The six detectors are retained as an interpretable, externally-validated decomposition (***Table S7***) and for setting per-mechanism thresholds (***Table S8***). The safety classifier was trained on 85 positives (80 FAIL_SAFETY, 5 FAIL_BOTH), of which 66 terminated in phase II against a phase III/IV PASS group, so it is power-limited and not phase-matched. Therefore, a phase III-restricted estimate was computed as the phase-matched comparison.

#### External validation of the safety axes

The cardiac and hepatic panels were validated against public sources (matched on IK14, ***Table S7***). For the cardiac axis, hERG channel-blockade labels^26^ (105 cohort compounds, 65 blockers) were predicted from the four cardiac toxicity-target binding features by five-fold cross-validated logistic regression (30 repeats), against a baseline of general promiscuity and lipophilicity. Organ-specificity was confirmed by residualizing the cardiac features on those confounds and bootstrapping over 2,000 compound resamples. Clinical concordance used SIDER^27^ counts of MedDRA cardiac-disorders terms (n = 438), with hepatic adverse-event counts as a specificity control. For the hepatic axis, the hepatic binding panel was tested against drug-induced-liver-injury DILIrank^28^ concern (n = 213). Each safety failure confirmed by the label audit was assigned an organ liability, classified from its audit rationale (***Table S9***).

#### Signal decomposition

Features were added in three cumulative steps, and a classifier using the same algorithm for that task was refit at each step. The first step used the disease/trial binary flags alone, the second added the within-fold disease complexity feature, and the third added the molecular-mechanism profile through the same per-fold top-20 selection. Each step’s marginal contribution is the paired fold-averaged AUROC increment over the preceding model.

#### Temporal (leave-future-out) validation

Training included trials initiated before a specified cutoff year, while testing included trials initiated in or after that year. Compounds present in training were excluded from the testing, ensuring that all test trials were both temporally future and structurally novel. Trials that started before the cutoff were used for training even when their outcome was reported after it.. Median imputation, the per-fold top-20 feature selection and within-fold disease complexity feature were all fit on the pre-cutoff training dataset, so no fit quantity used any post-cutoff trial. The molecular-mechanism profiles were extracted from a single current snapshot of public databases rather than from historically-frozen releases.

#### As-of-date *disease complexity* sensitivity analysis

To assess whether the disease complexity feature incorporates information from trials conducted after the trial being predicted, each indication’s failure rate was recalculated using only training-fold trials initiated in earlier years. Bayesian smoothing was retained, and the global rate was used for indications with no prior trials. Both feature definitions were fully evaluated while maintaining other parameters constant, and the results are reported alongside the signal decomposition.

#### structure-holdoutLabel audit

Every terminated-trial label included in the failure set (any of the FAIL classifications) was assessed by a blind, web-verified protocol of each trial’s ClinicalTrials.gov record and classified whether the termination was a drug-attributable safety or efficacy failure versus a non-drug stop. The resulting noise rate was balanced across MERIT’s confidence strata. The cumulative audit over all label-correction excluded 187 trials in total, comprising 134 non-drug stops, 47 mis-scoped outcomes and 6 with no posted results (***Table S10).*** A second annotator independently re-classified a blinded, stratified sample of 68 terminated-trial labels under the same frozen rubric, giving Cohen’s κ = 0.70 on the true-failure versus non-drug stop decision, with 86.6% raw agreement and a full six-category κ was 0.78.

#### Bootstrap stability and confidence intervals

AUROC confidence intervals (CI) were obtained by a compound-clustered bootstrap that resampled drugs (by SMILES) with replacement over 2000 resamples and recomputed the same fold-averaged AUROC as the final metric with corresponding 95% CI. Principal component analysis (PCA) stability was assessed via 100 drug-resamples (***Note S4***), and per-feature Cliff’s d CI via trial-level resampling over 2000 resamples.

#### Phase-matched efficacy analysis

Since PASS requires phase III completion while terminated efficacy failures span phases, trial phase is correlated with the efficacy label. The decomposition was repeated restricted to phase III trials only (2,114 PASS, 326 FAIL_EFFICACY), where both classes are present, under the identical structure-holdout protocol.

### Repositioning analysis

The repositioning analyses used MERIT’s efficacy classifier to score mechanism-fit. The out-of-fold predicted failure probability for a drug-indication pair, averaged over 5 seeds, defines a mechanism-fit score [fit = 1 − P(fail)]. The fit never use the drug’s identity or any trial outcome. The target-disease biology features exclude the Open Targets known-drug and literature channels, and a mechanism-fit variant with the disease base rate removed was evaluated as a sensitivity analysis.

#### Within-drug indication selection

For drugs with both a PASS and a FAIL_EFFICACY indication, the mechanism-fit was compared between the two indications for the same drug using a paired Wilcoxon test. Comparing within the same drug removes cross-drug confounds.

#### Recovery of known repositionings

Cohort compounds that failed one indication for efficacy but are approved for another in repoDB^29^, matched by drug name, define a known repositioning set of 60 compounds. Recovery is the fraction with the approved indication scored above the failed one. This set is preselected on having a known successful repositioning, so the recovery fraction is a within-pair ranking accuracy and is not generalizable to all failed drugs.

#### Novel nominations

For compounds that failed for efficacy with no approved indication, target genes were resolved through ChEMBL and scored against all diseases by Open Targets genetic association. The highest-supported indications a compound was not trialed for are reported as hypotheses. Each nominated indication is checked against Orphanet^30^ and MONDO^31^ for rare disease classification. Nominations are pre-trial mechanism hypotheses, not efficacy predictions.

#### Mechanism-fit validation on later indications

The efficacy classifier was retrained only on trials that were initiated before a cutoff year (2016/2017/2018) and used to score mechanism-fit for compounds with a pre-cutoff efficacy failure and no completed phase III/IV trial before the cutoff against their post-cutoff new indications (***Table S11***). Metrics are computed for each drug-indication nomination, with COVID/coronavirus conditions grouped into one indication.

### Prospective registration

To prospectively evaluate the model, predictions were fixed for ongoing trials before outcomes are reported. All ongoing phase III trial, including those involving the cohort compounds and the novel drug-indication pairs, were retrieved from the ClinicalTrials.gov v2 API. Trials listing multiple conditions, combination treatments, or sub-studies, and trials meeting the cohort’s efficacy exclusion criteria were removed. The 207 drug-level molecular features were reused from the compound, as well as the 10 indication-context features. The target-disease mechanism-fit was recalculated for each drug-indication pair using public biological data, while trial-context features were derived from each trial’s protocol. No median imputation was applied. The overall PASS-versus-failure classifier, trained on the full modeling cohort, generated calibrated failure probabilities. After applying novelty, combination-attribution and outcome-blindness filters, 55 novel predictions remained, spanning 54 trials and 37 compounds. All 55 predictions constitute the primary registered evaluation set, with no post-hoc selection, and will be assessed at readout using AUROC and calibration. Each prediction was also assigned a pre-registered confidence category of high-confidence or uncertain (***Note S5***). The complete prediction set, including calibrated failure probabilities, SHA-256 hash and generating commit, was deposited at https://doi.org/10.5281/zenodo.21824277 and https://github.com/OmicMD/MERIT, and before any trial outcomes were reported.

### Transcriptomic perturbation analysis (evaluated, not adopted)

LINCS L1000 consensus perturbation signatures^32^ were used to test whether drug-induced transcriptional response improves prediction in addition to the structure-, network- and tissue-derived features. These signatures were linked to compound structures via the Drug Repurposing Hub^33^ and matched on IK14 (446 profiled compounds). All analyses were evaluated within the jointly profiled subset, since signature availability is itself outcome-correlated. Two methods were tested. The first was connectivity reversal, which scores a drug by how different its gene-expression signature is to the disease’s own signature. The second method tested for a network connection between the drug’s perturbed genes and the disease’s associated genes, using the STRING and OmniPath^34^ protein networks. Both were scored against four nulls across a range of seed thresholds and restart rates. No transcriptomic feature was incorporated into MERIT.

## RESULTS

### MERIT integrates biological profiling of small-molecule compounds

This study assembled 3,133 trials records (2,967 distinct trials and 2,200 drug-indication pairs) across 753 small-molecule compounds from ClinicalTrials.gov^10^, classified as PASS, FAIL_EFFICACY, FAIL_SAFETY or FAIL_BOTH by a manual audit of every terminated trial (***Figure 1a***). PASS denotes phase III/IV completion, not approval or demonstrated efficacy, a permissive label confounded with trial phase. Restricting evaluation to phase III trials had little effect on AUROC (efficacy changed from 0.765 to 0.753, and overall changed from 0.770 to 0.748), indicating that the model distinguishes trial outcomes rather than relying on differences between trial phases. Each compound was profiled with eight compound-level pre-trial modules covering tissue-specific i (Human Protein Atlas^14^), pathway engagement (STRING^15^ against Open Targets^16^ disease genes), binding specificity, safety pharmacology, pharmacokinetics (DruMAP^17^), target-disease biology, genetics support^16, 35^, indication and trial context, disease complexity, with a ninth trial-design module added during cross-validation (***Figure 1b***). Each of the five structure-computed modules alone achieved AUROC of only 0.682 to 0.707 (***Figure S1***), indicating that the predictive accuracy of MERIT is due to the combination of the modules.

**Figure 1.**
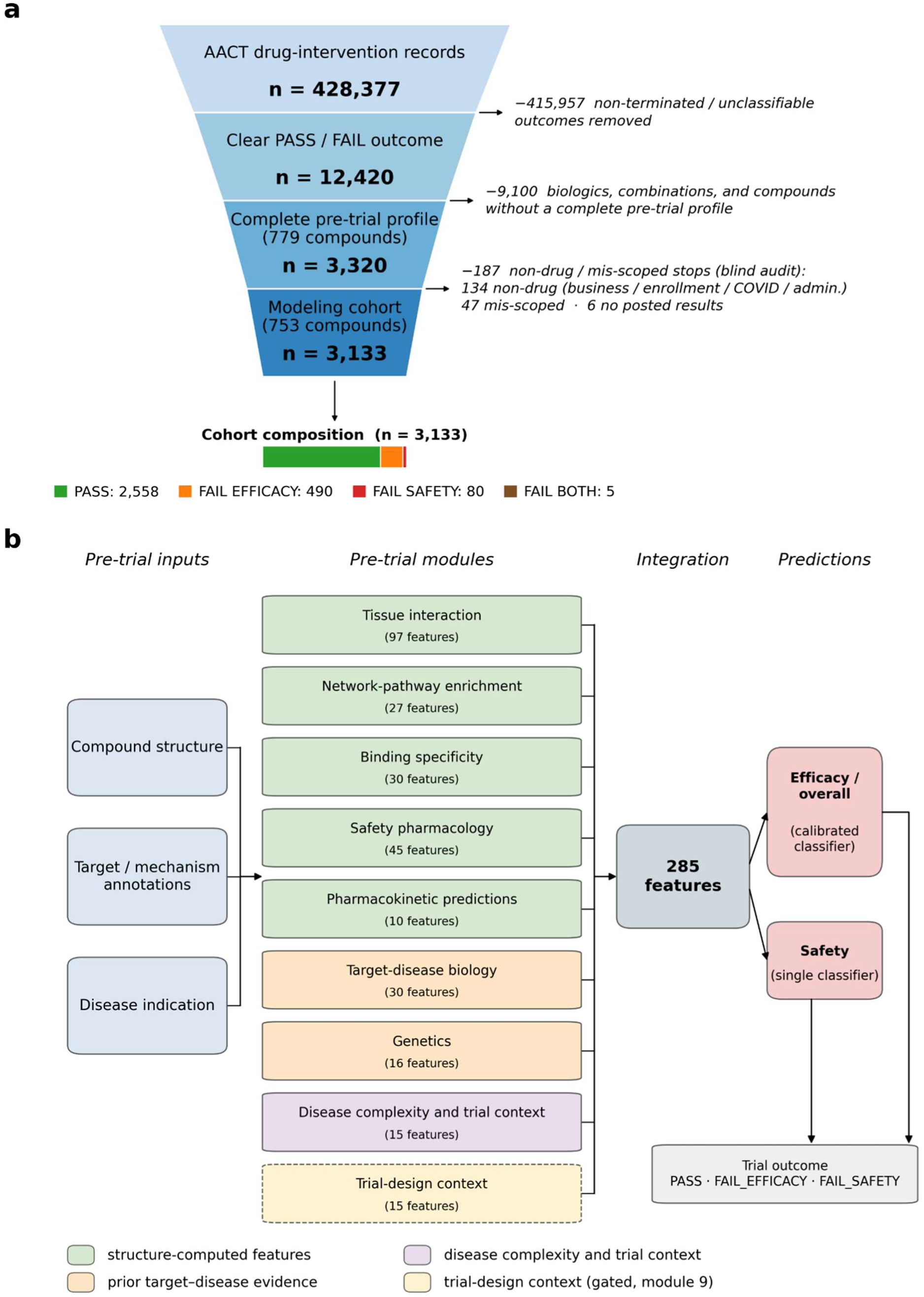
Dataset and computational architecture. No single data source predicts trial outcome on its own, and MERIT predicts from a combination of eight modules. **(a)** Trial-selection funnel: 428,377 AACT records narrow to the modeling cohort of 3,133 trials across 753 compounds after the blind label audit. **(b)** MERIT computational architecture: eight core pre-trial modules plus a gated ninth trial-design module, combined into 285 features and routed to two classifiers (one for efficacy or overall, and a single classifier over the six safety axes).

### MERIT discriminates outcomes for structurally unseen drugs

Tested only on drugs whose structure was not in training^9^, the full model reaches an AUROC of 0.770 overall, 0.765 for efficacy, and 0.784 for safety (***Figure 2a***). All drug-clustered 95% CIs excluded chance (0.745 to 0.796 overall, 0.742 to 0.790 for efficacy, and 0.737 to 0.831 for safety), while shuffling outcome labels reduced AUROC to 0.498 across 50 permutations (***Figure 2b, Table S12***). Among the 100 trials assigned the highest overall risk, the failure rates were four to six times the cohort baseline of 18.4% (***Table S13***).

**Figure 2.**
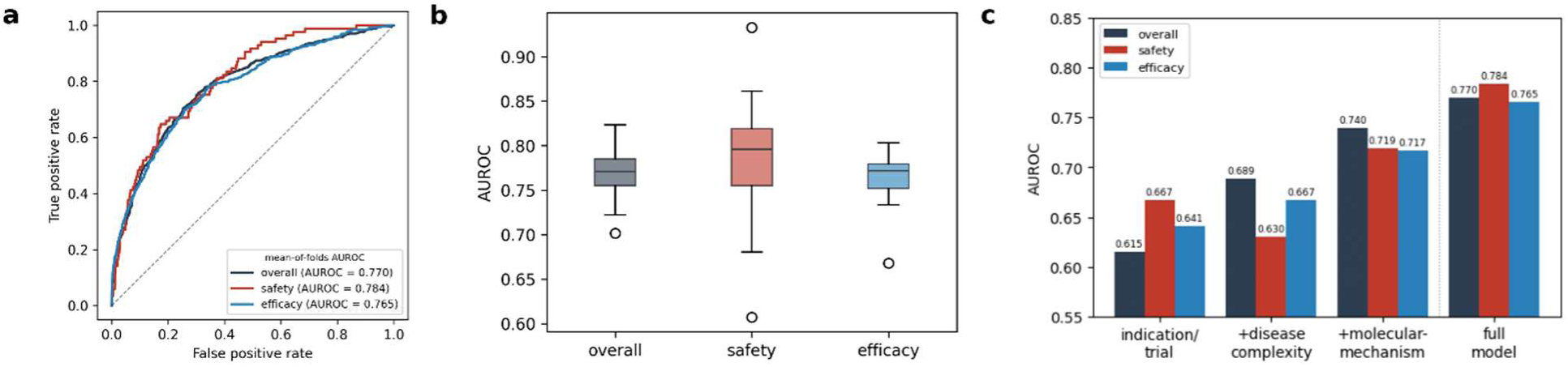
Prediction performance and signal decomposition. MERIT differentiates between trial outcomes for novel drugs, and the molecular-mechanism profiles account for most of the efficacy increment. **(a)** AUROC curves for the full model under structure-holdout. Overall AUROC is 0.770, safety is 0.784 and efficacy is 0.765. **(b)** Per-fold AUROC across the 25 folds (5 seeds × 5 folds). **(c)** Signal decomposition (averaged across folds) starting from indication and trial flags, adding the disease complexity baseline and then the molecular-mechanism profile. Molecular-mechanism profile contributes most to the efficacy AUROC prediction (***Table S12***).

AUROC measures how well the model ranks trials by risk, but does not report performance at a specific classification threshold (Table S4). Accuracy alone can be misleading because failures are uncommon (18.4% overall, 17.5% for efficacy, and 3.2% for safety). Predicting success for every trial would achieve 81.6% overall accuracy while missing every failure. At the default 0.50 threshold, the model identified failures with high precision (0.729) but low recall (0.118). Lowering the threshold to each task’s failure base rate resulted in better balanced, identifying 71.7% of overall failures with 34.6% precision (F1 of 0.449 and balanced accuracy of 0.692). Efficacy showed similar performance (F1 of 0.419 and balanced accuracy of 0.678). Safety predictions were less precise (0.144), reflecting the small number of observed failures (n = 85) relative to the 20 selected features. PR-AUC exceeded the corresponding failure base rates several fold across all three tasks (0.434 overall, 0.422 for efficacy and 0.226 for safety). Full results, including uncertainty estimates, MCC, threshold-dependent metrics, are reported in ***Table S4***.

### MERIT outperforms published methods on the same trials

MERIT was evaluated against TrialBench^7^ and HINT^6^ on identical trials and structure-holdout folds. These methods were selected because both provide publicly available code, support small-molecule drug-indication pairs, and can be evaluated on our cohort. They also represent distinct approaches. HINT uses a deep learning model over molecular graphs, disease codes and eligibility text (HINT), whereas TrialBench uses tabular features spanning multiple aspects of trial design.

Each model was evaluated under two conditions, a strict comparison excluding enrollment count and establishment metadata from all models, and a comparison retaining these features under each method’s original configuration. MERIT outperformed both methods under both conditions, with higher AUROC by 0.131 over TrialBench and 0.078 over HINT (***Table 1, Figure S2***).

**Table 1.** Comparison with published trial-outcome methods on shared trials.

| Model | Task | Reported | Strict | MERIT | $\Delta(\text{MERIT-strict})$ | *P* |
| --- | --- | --- | --- | --- | --- | --- |
| HINT | phase III completion | 0.654 | 0.626 | 0.704 | +0.078 | paired t-test $7 \times 10^{-9}$,<br>Wilcoxon $2 \times 10^{-7}$,<br>DeLong $3 \times 10^{-9}$ |
| TrialBench | Drug approval | 0.794 | 0.637 | 0.768 | +0.131 | paired t-test $1 \times 10^{-6}$,<br>Wilcoxon $6 \times 10^{-5}$ |

- 2 Each model was evaluated using its original outcome definition (phase III completion for HINT and drug approval for TrialBench). 3 The comparisons assess distinct prediction tasks and are not directly comparable.

On the 1,142 shared trials comprising 412 compounds, TrialBench achieved and AUROC of 0.794 using its full feature set. Removing enrollment count and establishment metadata reduced AUROC to 0.637 (***Table 1***). On contrast, MERIT’s mechanism-based features achieved an AUROC of 0.768, outperforming TrialBench’s trial-design features by 0.131 across 15 folds (paired *t*-test *P* = 1×10^-6^; ***Table 1, Figure S2a***). Combining both feature sets further increased AUROC to 0.807.

On shared phase III cohort of 2,787 trials, HINT was retrained and evaluated using structure-holdout evaluation. MERIT achieved an AUROC of 0.704 against 0.626 for HINT, in 24 of 25 splits (paired *t*-test P = 7×10^-9^; ***Table 1, Figure S2b***). Both models performed better under HINT’s structure-blind evaluation, with MERIT achieving an AUROC of 0.737 against HINT’s 0.654.

Other published methods were less directly comparable. Since inClinico^5^, released neither its model nor feature set, an approximation model for its phase II-to-III transition task (***Note S6, Figure S3***) was reconstructed. Lo et al.^8^ and Feijoo et al.^36^ predict approval from sponsor and registry variables without molecular features, while knowledge-graph models such as TxGNN^37^ predict drug-indication plausibility, not the individual trial outcomes. A classical network-proximity predictor^25^ was also evaluated and scored efficacy at chance (AUROC 0.470, 95% CI of 0.406 to 0.538).

#### Predictive performance is not specific to the learning algorithm

Random forest, logistic regression and single gradient-boosting algorithm were compared on all three prediction tasks using identical features (***Table S6***). Random forest outperformed logistic regression by 0.060 overall and 0.076 for efficacy. Random forest slightly outperformed gradient-boosting across all three tasks (AUROC differences of +0.028 overall, +0.019 for efficacy and +0.022 for safety). MERIT’s final efficacy AUROC of 0.765 exceeded the single algorithm AUROC of 0.722 because it combines prediction from three gradient-boosting implementations. Predictive performance comes from the features rather than the classifier.

### Mechanism-fit features contributes more than disease complexity

The contribution of each module was measured by sequential addition of features using identical training and test datasets. (***Figure 2c, Table S12***). For overall trial outcomes, indication and trial context achieved an overall AUROC of 0.615. Adding disease-complexity, calculated exclusively from training data, increased AUROC to 0.689, while molecular-mechanism profile increase it further to 0.742. The molecular-mechanism profile contributed more to efficacy prediction than disease complexity, improving AUROC by 0.051 versus 0.026. The molecular-mechanism profile alone achieved an efficacy AUROC of 0.695, exceeding the 0.666 obtained from indication and trial context combined with disease complexity. These results identify molecular-mechanism profile as the better predictor of efficacy outcomes.

Efficacy prediction depends on matching a drug’s mechanism to the disease it is intended to treat. The improvement in AUROC comes from features that measure this match, including genetic and Mendelian evidence, Open Targets biology evidence, target-disease network connections, KEGG pathway membership, and DepMap dependency. This contribution persisted when analysis was restricted to phase III trials (+0.054 AUROC on 2,114 completed trials and 326 efficacy failures, compared to +0.051 in the full cohort).

Safety prediction differs because failure can occur due to any of several distinct liabilities. These are grouped into six safety detectors consisting of promiscuity, hepatic, cardiac, network, tissue, and trial context. After removing shared disease complexity component, the six detectors’ top-decile alerts were combined and this identified 26% of safety failures, compared to 18% for the dominant individual detector (***Note S7***). A joint classifier combining all six detectors outperformed a noisy-OR model that treats each liability independently, achieving an AUROC of 0.784 versus 0.725 (higher in 23 of 25 folds, *P* < 10^-5^). The joint classifier was therefore retained, with the individual detectors providing a decomposition of its safety predictions (***Table S7***).

External datasets provided independent validation of the six safety detectors. The cardiac detector was associated with *in vitro* hERG blockade and clinical cardiac events in SIDER. In contrast, hepatic detector showed no association with DILIrank liver-injury classification, potentially reflecting the influence of patient-specific factors on drug-induced liver injury (DILI; ***Tables S7, S9*** & ***Note S8***).

#### The safety task is power-limited and not phase-matched

The safety classifier was trained on 85 positives (80 FAIL_SAFETY and 5 FAIL_BOTH) against 2,555 completed trials. Its performance was less stable than the overall classifier, with a wider drug-clustered 95% CI (0.737 to 0.831 versus 0.745 to 0.796) and larger variability across folds (SD of 0.066 versus 0.026).

Safety failures and completed trials also differed substantially by phase. Most safety failures occurred in phase II (66 of 85), whereas completed trials required phase III/IV completion. Restricting both groups to phase III (including phase II/III) reduced safety AUROC from 0.784 to 0.731, a larger decrease than for efficacy (0.765 to 0.753) or overall (0.770 to 0.748) predictions. The phase-matched AUROC of 0.731, based on only 19 safety failures, is more relevant for phase III decisions.

These limitations make the safety classifier preliminary and hypothesis-generating, rather than suitable for clinical deployment or direct comparison with published safety predictors. Adding 32 independently audited safety failures did not improve prediction, potentially because the additional cases involved patient-specific HLA-dependent mechanisms (***Note S9***). External hERG, SIDER and DILIrank analyses provide supporting evidence for individual safety signals, but do not resolve the limitations of the trial outcome classifier.

Finally, the feature contributions reported here establish predictive associations, not causal mechanisms. Genetic support and target-disease relationships may help distinguish successful from unsuccessful trials, but do not establish that engaging a particular target causes clinical benefit. MERIT provides *mechanism-associated prediction*, which uses biological evidence connecting drug targets to diseases for predicting trial outcomes. Establishing *mechanistic explanation*, in which a specific molecular event is shown to cause an individual trial, requires experimental validation beyond the scope of this study.

### Temporal validation on structurally novel compounds

Trained on trials initiated before a specified cutoff year, and tested on trials initiated in or after that year with every training compound removed, MERIT achieved an AUROC of 0.704 to 0.737 overall and 0.649 to 0.731 for efficacy across 2016 - 2018 cutoffs, with safety based on only 23 to 31 positives. These are lower than the full-cohort overall and efficacy AUROC (0.770 and 0.765) because each temporal split trained on fewer trials (***Table S14***).

#### The disease complexity feature is robust to as-of-date restriction

An indication’s historical failure rate over the whole dataset would incorporate outcomes of trials that were conducted after the trial being predicted. This was prevented by computing this feature strictly inside each cross-validation training fold and it was applied on the pre-cutoff training dataset only. Neither prevents a trial from training that started later from contributing to an earlier test trial’s value.

The feature was rebuilt as of each trial’s start date, and indications without prior trials used the global rate. Under this strictly-prior encoding with remaining factors maintained constant across 25 folds, the overall AUROC changed from 0.739 to 0.731, and efficacy from 0.722 to 0.713 (single classifier configuration, ***Table S12***). Both changes are negligible and within the fold-to-fold SD (0.026 - 0.037), indicating that the as-of-date restriction does not reduce the contribution by the disease complexity feature. MERIT retains the within-fold encoding so the model matches the released feature matrix and predictions.

#### Historical-snapshot limitation of the biological databases

The temporal analysis split the trials at a historical cutoff year but derived the biological features from current releases of Open Targets, STRING, ClinGen/ClinVar, KEGG and DepMap. Current releases contain evidence published after the cutoff, so the analysis scores each trial using knowledge that was not available when the trial began. Archived releases of these resources are published, but they are versioned by release date rather than by evidence date, and Open Targets association scores are recomputed at each release against the then-current evidence base, so an archived release does not represent what was known at a given cutoff.

Two considerations limit how much this increases the temporal AUROCs, without removing the limitation. The target-disease biology and genetic features are dominated by genetic association evidence and network enrichment, which accumulate from disease-genetics studies, so post-cutoff additions are indirectly related to the trial outcome predictions. The temporal AUROCs remain an estimate of historical prospective predictions. In contrast, the locked prospective registration of 55 ongoing phase III drug-indication pairs does not depend on archived database releases.

### Population and endpoint mismatch precede efficacy failure

A trial outcome is dependent on the enrolled population and the primary endpoint measured. A drug with high mechanism-fit score can fail due to either of these factors and both can be assessed before the trial begins. A targeted drug fails more often when patients are not selected for that target’s role in the indication (70% of 10 trials without biomarker selection versus 15% in biomarker-selected patients). In a multi-system indication, a surrogate endpoint may be involved in different physiological processes than the one for the drug’s target leading to failure. Across six organ systems, 60% of trials with a mismatched endpoint failed against 9% of trials with a matched endpoint. For example, empagliflozin failed with a six-minute-walk endpoint (EMPERIAL^38^) that is not affected by its renal and fluid-handling mechanism, while succeeding on cardiovascular outcomes (EMPEROR-Reduced^39^). Tolvaptan succeeded on a kidney-volume endpoint, a process controlled by its target (TEMPO 3:4^40^). Three outcome-blind features in module 9 encode these population and endpoint mismatches (***Note S1***). Adding these features changed the risk ranking for the individual trials but the overall AUROC of 0.770 remained unchanged, suggesting that these features provide explanations for individual mismatch cases only. The remaining prediction errors are trials with a high mechanism-fit score and a matched endpoint but the trial failed because the treatment effect was too small.

### Misclassifications fall into two groups

There are two groups of high-confident misclassifications (***Figure 3a***). The safety misclassifications are drugs managed by dose control (e.g. taxanes, PARP and multi-kinase inhibitors), including known side-effects or patient-specific effects (***Table S15***). The efficacy misclassifications are approved drugs that failed in a particular population, formulation or line of therapy (e.g. lacosamide and mesalamine). These failures depend on the treatment effect size. MERIT detects risks and mechanisms within a characterized boundary rather than predicting failure in general. The per-drug feature profiles in ***Figure 3a*** are post-hoc interpretations of individual cases, not validated causal attributions.

**Figure 3.**
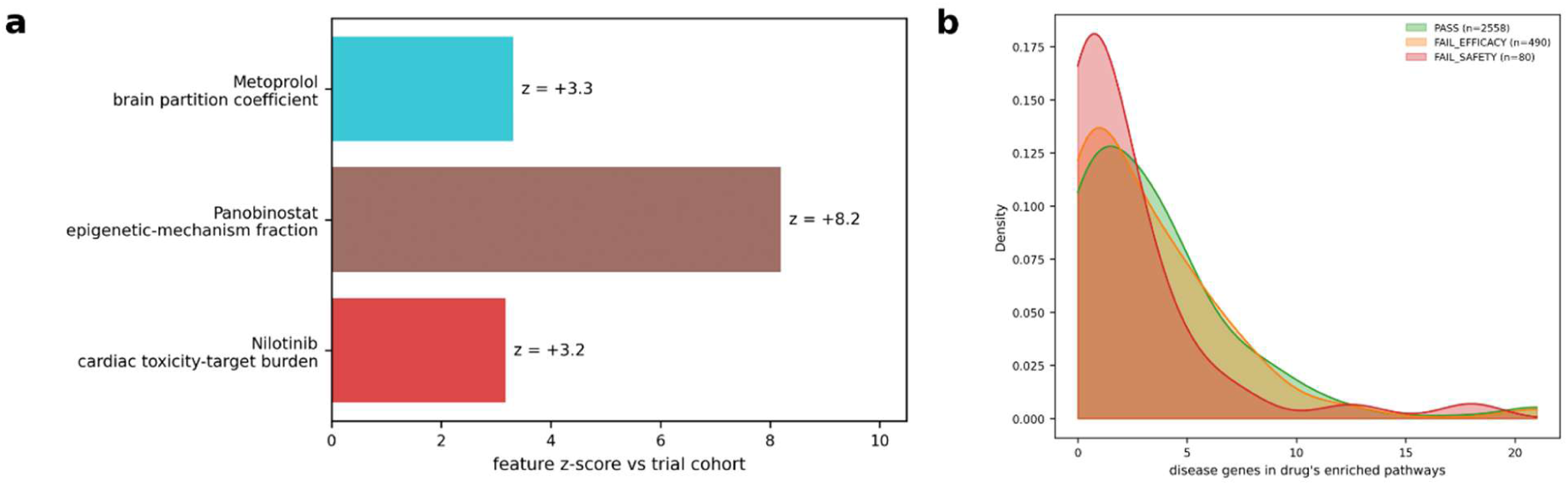
Confident misclassifications and target-disease pathway overlap. One extreme feature explains individual misclassifications without increasing overall accuracy, and safety failures drugs bind targets outside the disease’s gene network. **(a)** Three misclassifications, each with the one feature deviating most from the cohort mean (z-score). Nilotinib’s feature is cardiac-target burden (z = +3.2), panobinostat’s is the epigenetic-mechanism fraction (z = +8.2), and metoprolol’s is the brain-partition mismatch against chronic obstructive pulmonary disease (COPD, z = +3.3). **(b)** Disease-pathway overlap. Overlap counts the disease’s associated genes that are found in the drug’s targets associated pathways. Curves are kernel density estimates over trials. Safety failures have the lowest overlap (median 1, versus 2 for efficacy failures and 3 for passes), consistent with off-target toxicity.

A drug-indication pair tested in multiple trials can have different outcomes, where drug-indication pairs that failed once passed in another. This was observed in 18% of drug-indication pairs (95% CI of 14 to 22%). Dose, population, endpoint and chance contribute partially to efficacy outcome. None of these is predictable from pre-trial biology features since a model that uses only these features assigns one score to all trials of that drug-indication pair. Among the confident efficacy misclassifications, 23% of the cohort contains a completed trial of the same drug-indication pair (***Note S9***). An efficacy AUROC of 0.765 reflects the limitations of pre-trial biology features on this cohort.

Three features differentiated safety failures from passes (Cliff’s d = -0.25, 2000-resample CI excludes zero). These are disease-pathway overlap, the fraction of the drug’s enriched pathways that contain disease genes, and the number of disease genes in those pathways (***Note S4***). These features provide an explanation for the different safety outcomes (***Figure 3b***). Efficacy outcomes differed little when using these features (d = -0.05).

### Repositioning failed drugs by mechanism-fit score

The same mechanism-fit score can be used to generate hypotheses since the score is computed using structure-based and target-disease features (***Figure 4***).

**Figure 4.**
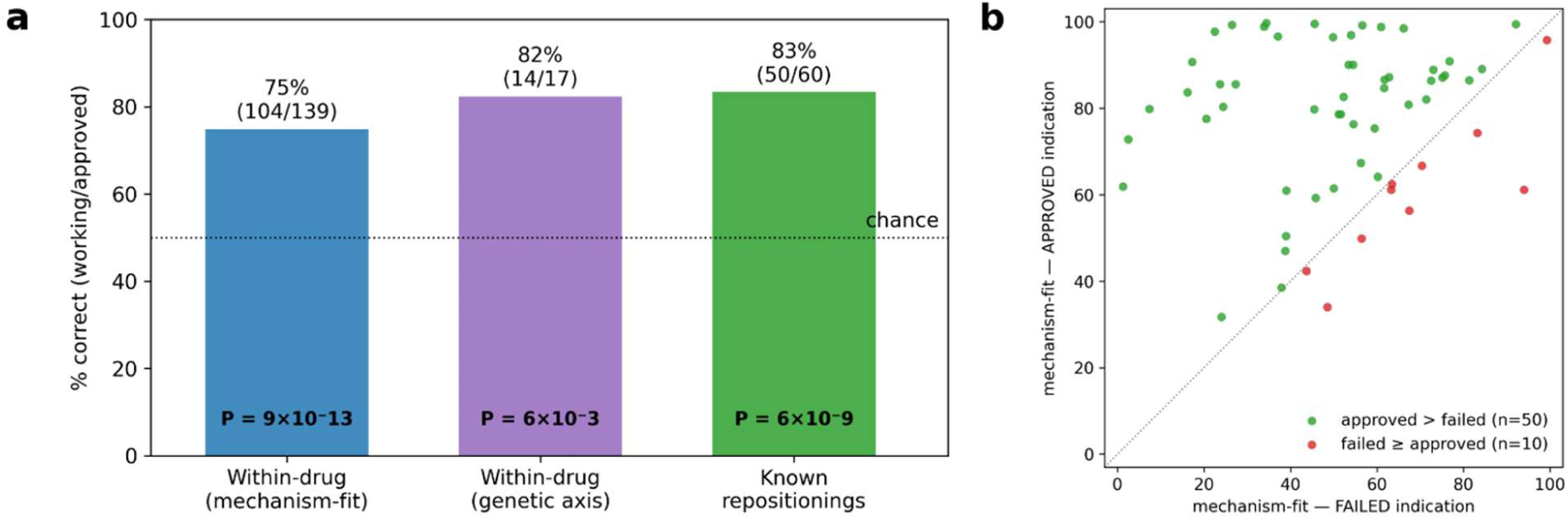
Repositioning failed drugs. MERIT ranks a drug’s working indication above its failed one by mechanism-fit score. **(a)** The working indication had higher mechanism-fit score in 75% of drugs (*P* = 9×10^-13^), and higher genetic association score in 82% of informative drugs (*P* = 6×10^-3^). MERIT ranked the approved indication above the failed one in 83% of known repositionings from repoDB (*P* = 6×10^-9^). **(b)** For the 60 drugs with known repositionings, the mechanism-fit mean score for the approved indication is 0.779 versus 0.514 for failed indications (paired Wilcoxon *P* = 6×10^-9^).

Across 139 compounds with both a passing and an efficacy-failing indication, the mechanism-fit score was higher for the working indication in 104 compounds (75%, *P* = 9×10^-13^; ***Figure 4a***). The genetic association ranked the working indication higher in 14 of the 17 compounds (*P* = 6×10^-3^). Across the 60 cohort compounds that failed one indication for efficacy but are approved for another in repoDB, MERIT scored the approved indication above the failed one in 50 cases (mean mechanism-fit of 0.779 versus 0.514, *P* = 6×10^-9^; ***Figure 4b***). Recovered compounds include tamoxifen, which scored higher for breast cancer than for Duchenne muscular dystrophy, and atorvastatin, which scored higher for hypercholesterolemia than for bicuspid aortic valve (***Table S16***). This set of 60 compounds was preselected so the 83% represents the fraction of drug-indication pairs that were correctly ranked. It does not imply that 83% of failed drugs can be repurposed. The mechanism-fit score used none of the Open Targets known-drug or literature channels. Under a sensitivity analysis, where each disease’s base rate was removed from the score, MERIT still ranked the approved indication above the failed one for 42 of the 60 compounds (70%). For example, MERIT scored chronic myeloid leukemia and gastrointestinal stromal tumors higher than the failed indications, COVID-19 and loiasis.

For 154 compounds that failed for efficacy and have no approved indication, MERIT nominated new indications (***Table S17***) for 52 compounds, consisting of 146 nominated drug-indication pairs. Published evidence showed that 129 pairs (88%) had clinical, target-class or preclinical support and 97 pairs (66%) had a registered trial of either the same drug or a same-target agent (***Table S18***). Galeterone, a CYP17A1 inhibitor that a failed phase III prostate cancer trial (ARMOR3-SV), was nominated for congenital adrenal hyperplasia, where another inhibitor abiraterone is now in trials (***Table S19***). These nominations are pre-trial mechanism hypotheses, not efficacy claims.

Finally, the efficacy classifier was tested on repositionings that were confirmed only after the training cutoff. Mechanism-fit scores for compounds with a post-cutoff trial that passed was higher than a trial that failed efficacy, at both the 2016 (AUROC of 0.781, *P* = 0.032) and 2017 cutoffs (AUROC of 0.759, *P* = 0.025; ***Table S11***). For example, selinexor scored highest for multiple myeloma at both cutoffs, and siponimod scored multiple sclerosis above the failures at the 2016 cutoff. Each cutoff has 8 to 12 confirmed successes, consistent with the main result. The exception is COVID-19, where host-directed compounds were scored above observed failure rate. These compounds act on immune targets, whereas the disease is caused by a virus, and no pre-cutoff trials existed for COVID-19 to establish that distinction.

## DISCUSSION

This study shows that a drug’s trial outcome can be predicted before the trial begins, using molecular and disease features computed without any information on the successes of similar compounds. Two pre-trial inputs accounted for most of the prediction, the disease complexity, and the match between the compound’s mechanism and that indication. The disease complexity increased the overall AUROC from 0.615 to 0.689, and the molecular-mechanism profile added a further 0.053. On the efficacy task, the molecular-mechanism profile increased AUROC by 0.051 and disease complexity added 0.026, with structure-holdout evaluation limiting both.

Efficacy failure was associated with a mismatch between the compound’s mechanism and the disease, whereas safety failure can occur from only one liability. A joint classifier over the six safety detectors achieved higher AUROC than any individual detector. Similarly, splitting the efficacy task into independent detectors reduced the prediction AUROC. Genetic analyses of why trials stopped supported these findings^41^. The mechanism-fit features also identified target-disease relationships without direct gene overlap, such as letrozole and azacitidine^35, 42^. Drug-induced expression changes did not improve prediction beyond MERIT’s features (446 LINCS L1000 compounds^32, 33^; ***Note S9***).

Knowledge-graph repositioning models score the co-occurrence of a drug-indication pair in both literature and curated databases^37^. MERIT scored each pair from mechanism-fit, using the same classifier that predicts efficacy failure. Ranking the approved indication above the failed one with the compound held out of training (***Figure 4***) shows that compounds which failed for efficacy can be screened against alternative indications.

MERIT’s overall AUROC of 0.770 is comparable to prior reports (AUROC of 0.70 to 0.85)^8, 36, 43^, but those did not use a structure-holdout evaluation. The highest, inClinico’s 0.882^5^ on phase II-to-III transition, used a split that held out neither compound nor target. Reproducing that task under a structure-holdout split resulted in AUROC of 0.746, mostly due to trial-registration details including sponsor, trial size, and a count of the program’s own later trials (***Figure S3; Note S6***). The structure-blind evaluation inflated the reported AUROC by 0.136, which is the reason structure-holdout evaluation should be the minimum standard.

The genetic association score is zero when the target or the disease is absent from Open Targets, and when the target is present without an association with the disease. The mechanism-fit features cannot distinguish missing evidence from evidence with no association. The disease complexity term increased the overall AUROC by 0.074 when it was the first module added to the indication and trial flags. However, when it was added to the model last, it only increased the overall AUROC by 0.014 (***Table S12***). The disease complexity relied on the indication’s historical failure rate, so in an indication that has usually succeeded, the model can underpredict failure. In the top quartile of indications with high historical failure rate, 45% of indications failed if the target is genetically associated with the disease, compared to 62% of those without genetic association. The failure rate is lower for compounds with genetic support for the target in these indications.

MERIT’s outcome label is trial completion at phase III/IV, taken from the registry, with terminated trials classified by blind audit as drug-attributable efficacy or safety failures or as non-drug stops. It predicts whether a trial completes rather than terminates for a drug-attributable reason. It does not predict primary endpoint success, since a trial that completes while missing its primary endpoint is labeled PASS. It also does not predict regulatory approval, which depends on effect size, comparator, regulatory judgment and commercial decisions outside the model’s features. Finally, it is not a measure of pharmacological efficacy.

The PASS classification consists of trials that met their primary endpoint, and completed trials that missed the primary endpoint. The efficacy task separates drug-attributable efficacy terminations from all other trials, not effective drugs from ineffective ones. A trial that completes without termination is labeled PASS, so the pass rate under this definition is higher than under an approval-based one. Hence, the TrialBench comparison was computed using TrialBench’s approval label. In the repositioning results, a high mechanism-fit score hypothesize that a mechanism suits an indication, not a prediction that a trial would meet its endpoint.

The safety classifier is preliminary and hypothesis generating, not a tool for development decision, because it was trained on only 85 safety-failure trials that are not phase-matched to the phase III/IV PASS group. The phase III-matched AUROC of 0.731 was based only on 19 safety failures (***Note S9***). The predicted safety-failure score was uncorrelated with black-box warning or post-market withdrawal (AUROC of 0.501 and 0.483; ***Notes S9***), so it predicted trial termination risk specifically. Most confident safety misclassifications share target with other drugs from trials labeled PASS. A failure and its safe counterpart differed in dose, exposure and population, so an indication-specific threshold was applied for each risk tier by indication (***Table S8***). Restricting the cohort to small molecules is intentional, similar to structure-dependent models such as HINT.

All reported AUROCs are retrospective, so we registered locked predictions for 55 novel drug-indication pairs across 54 ongoing phase III trials (mechanism recomputed per pairing). Each prediction has a calibrated failure probability fixed by a public timestamp and SHA-256 hash (deposited at https://doi.org/10.5281/zenodo.21824277). The entire set will be scored against the trials’ actual outcomes, by AUROC and calibration. Each prediction was assigned to high-confidence or uncertain group, with the hypothesis that the 27 high-confidence pairs (24 PASS and 3 FAIL) is more accurate than the uncertain group. Predicting for structurally novel compounds, including compounds entering phase I, is the definitive prospective test.

Drug-attributable termination can therefore be predicted before enrollment begins, in time to change the compound’s indication before incurring high costs. A drug that fails its trial has not necessarily failed its biology, but may have been directed at the wrong disease. The same pre-trial features that anticipates the failure can nominate a better-matched indication.

## ASSOCIATED CONTENT

## Data and Software Availability

The clinical-trial data underlying this study are publicly available from the Aggregate Analysis of ClinicalTrials.gov (AACT). Molecular and biological annotations were derived from public resources (PubChem, ChEMBL, Human Protein Atlas, STRING, Open Targets, DruMAP, SIDER, DILIrank, repoDB and LINCS L1000). All analysis code is available at https://github.com/OmicMD/MERIT. The structure-activity-tissue binding profiles are produced by an unreleased proprietary pipeline. To keep the study reproducible, the pipeline’s frozen per-compound predicted binding profiles [∼3Tb of data] are available upon request, and all features derived from them are released in the cohort files, so model training and evaluation can be reproduced from the released data without access to the proprietary pipeline. Only generating binding profiles for new compounds requires the proprietary pipeline, which is available from the authors for non-commercial academic use upon request.

The processed trial- and arm-level modeling cohorts, the curated outcome labels with per-label provenance, the frozen per-compound pre-trial biology feature matrix and the per-fold out-of-fold predictions of the final model and benchmark comparisons are released at https://github.com/OmicMD/MERIT and https://doi.org/10.5281/zenodo.21824276. The locked prospective-registration predictions across the 55 ongoing phase III trials of novel drug-indication pairs, with their SHA-256 commitment are deposited at https://doi.org/10.5281/zenodo.21824277.

## Supporting Information

Tables S1 & S2 – Efficacy-evaluation exclusions, arm-level robustness (PDF)

Note S1, Figure S1 – Module features enumerations (PDF)

Note S2 – Target-disease mechanism axis (PDF)

Tables S4-S6 – Classification metrics, calibration, baseline models (PDF)

Note S3, Figure S2 – Benchmark reproduction protocol (PDF)

Note S4 – Bootstrap stability of the six molecular axes (PDF)

Note S5 – Confidence gate bands (PDF)

Note S6, Figure S3 – Reconstruction of a published phase II-III transition model (PDF)

Notes S7 & S8, Tables S7 & S9 – Safety mechanism detectors, external validation (PDF)

Table S12 – Signal decomposition, as-of-date disease complexity (PDF)

Tables S13-S15 – Precision at top-k, temporal validation, drug-class performance (PDF)

Table S10 – Terminated trial label audit (PDF)

Tables S11, S16-S19 – Repositioning validation and nominations (PDF)

Note S9, Tables S3 & S8 – Confident misclassification tests, safety decision layer (PDF)

Table S20 – Analysis scripts (PDF)

## Author Contributions

GR conceived and designed the study. CKT designed the binding profile pipeline. IM and BAS implemented and optimized the pipeline. SM managed the data and analysis infrastructure. All authors were involved in the analyses of data. CKT and GR wrote the manuscript draft, and all authors contributed to the manuscript revisions.

## References

(1) Hay, M.; Thomas, D. W.; Craighead, J. L.; Economides, C.; Rosenthal, J. Clinical development success rates for investigational drugs. Nat Biotechnol 2014, 32 (1), 40–51. DOI: 10.1038/nbt.2786. Wong, C. H.; Siah, K. W.; Lo, A. W. Estimation of clinical trial success rates and related parameters. Biostatistics 2019, 20 (2), 273–286. DOI: 10.1093/biostatistics/kxx069.

(2) Paul, S. M.; Mytelka, D. S.; Dunwiddie, C. T.; Persinger, C. C.; Munos, B. H.; Lindborg, S. R.; Schacht, A. L. How to improve R&D productivity: the pharmaceutical industry’s grand challenge. Nat Rev Drug Discov 2010, 9 (3), 203–214. DOI: 10.1038/nrd3078.

(3) Hwang, T. J.; Carpenter, D.; Lauffenburger, J. C.; Wang, B.; Franklin, J. M.; Kesselheim, A. S. Failure of Investigational Drugs in Late-Stage Clinical Development and Publication of Trial Results. JAMA Intern Med 2016, 176 (12), 1826–1833. DOI: 10.1001/jamainternmed.2016.6008.

(4) DiMasi, J. A.; Grabowski, H. G.; Hansen, R. W. Innovation in the pharmaceutical industry: New estimates of R&D costs. J Health Econ 2016, 47, 20–33. DOI: 10.1016/j.jhealeco.2016.01.012.

(5) Aliper, A.; Kudrin, R.; Polykovskiy, D.; Kamya, P.; Tutubalina, E.; Chen, S.; Ren, F.; Zhavoronkov, A. Prediction of Clinical Trials Outcomes Based on Target Choice and Clinical Trial Design with Multi-Modal Artificial Intelligence. Clin Pharmacol Ther 2023, 114 (5), 972–980. DOI: 10.1002/cpt.3008.

(6) Fu, T.; Huang, K.; Xiao, C.; Glass, L. M.; Sun, J. HINT: Hierarchical interaction network for clinical-trial-outcome predictions. Patterns (N Y) 2022, 3 (4), 100445. DOI: 10.1016/j.patter.2022.100445.

(7) Chen, J.; Hu, Y.; Cai, M.; Lu, Y.; Wang, Y.; Cao, X.; Lin, M.; Xu, H.; Wu, J.; Cao, X.; et al. TrialBench: Multi-Modal AI-Ready Datasets for Clinical Trial Prediction. Sci Data 2025, 12 (1), 1564. DOI: 10.1038/s41597-025-05680-8.

(8) Lo, A. W.; Siah, K. W.; Wong, C. H. Machine Learning with Statistical Imputation for Predicting Drug Approval. Harvard Data Science Review 2019. DOI: 10.1162/99608f92.5c5f0525.

(9) Vamathevan, J.; Clark, D.; Czodrowski, P.; Dunham, I.; Ferran, E.; Lee, G.; Li, B.; Madabhushi, A.; Shah, P.; Spitzer, M.;, et al. Applications of machine learning in drug discovery and development. Nat Rev Drug Discov 2019, 18 (6), 463–477. DOI: 10.1038/s41573-019-0024-5.

(10) Zarin, D. A. T., Tony; Williams, Rebecca J.; Califf, Robert M.; Ide, Nicholas C. The ClinicalTrials.gov Results Database — Update and Key Issues. N Engl J Med 2011, 364, 852–860.

(11) Kim, S.; Chen, J.; Cheng, T.; Gindulyte, A.; He, J.; He, S.; Li, Q.; Shoemaker, B. A.; Thiessen, P. A.; Yu, B.;, et al. PubChem 2023 update. Nucleic Acids Res 2023, 51 (D1), D1373–D1380. DOI: 10.1093/nar/gkac956.

(12) Zdrazil, B.; Felix, E.; Hunter, F.; Manners, E. J.; Blackshaw, J.; Corbett, S.; de Veij, M.; Ioannidis, H.; Lopez, D. M.; Mosquera, J. F.;, et al. The ChEMBL Database in 2023: a drug discovery platform spanning multiple bioactivity data types and time periods. Nucleic Acids Res 2024, 52 (D1), D1180–D1192. DOI: 10.1093/nar/gkad1004.

(13) Landrum, G. RDKit: Open-Source Cheminformatics Software. https://www.rdkit.org/ (accessed 2023.

(14) Uhlen, M.; Fagerberg, L.; Hallstrom, B. M.; Lindskog, C.; Oksvold, P.; Mardinoglu, A.; Sivertsson, A.; Kampf, C.; Sjostedt, E.; Asplund, A.;, et al. Proteomics. Tissue-based map of the human proteome. Science 2015, 347 (6220), 1260419. DOI: 10.1126/science.1260419.

(15) Szklarczyk, D.; Kirsch, R.; Koutrouli, M.; Nastou, K.; Mehryary, F.; Hachilif, R.; Gable, A. L.; Fang, T.; Doncheva, N. T.; Pyysalo, S.;, et al. The STRING database in 2023: protein-protein association networks and functional enrichment analyses for any sequenced genome of interest. Nucleic Acids Res 2023, 51 (D1), D638–D646. DOI: 10.1093/nar/gkac1000.

(16) Ochoa, D.; Hercules, A.; Carmona, M.; Suveges, D.; Baker, J.; Malangone, C.; Lopez, I.; Miranda, A.; Cruz-Castillo, C.; Fumis, L.;, et al. The next-generation Open Targets Platform: reimagined, redesigned, rebuilt. Nucleic Acids Res 2023, 51 (D1), D1353–D1359. DOI: 10.1093/nar/gkac1046.

(17) Kawashima, H.; Watanabe, R.; Esaki, T.; Kuroda, M.; Nagao, C.; Natsume-Kitatani, Y.; Ohashi, R.; Komura, H.; Mizuguchi, K. DruMAP: A Novel Drug Metabolism and Pharmacokinetics Analysis Platform. J Med Chem 2023, 66 (14), 9697–9709. DOI: 10.1021/acs.jmedchem.3c00481.

(18) Rehm, H. L. B., Jonathan S.; Brooks, Lisa D.; Bustamante, Carlos D.; Evans, James P.; Landrum, Melissa J.; Ledbetter, David H.; Maglott, Donna R.; Martin, Christa Lese; Nussbaum, Robert L.; Plon, Sharon E.; Ramos, Erin M.; Sherry, Stephen T.; Watson, Michael S. ClinGen — The Clinical Genome Resource. N Engl J Med 2015, 372, 2235–2242.

(19) Kanehisa, M.; Goto, S. KEGG: kyoto encyclopedia of genes and genomes. Nucleic Acids Res 2000, 28 (1), 27–30. DOI: 10.1093/nar/28.1.27.

(20) Tsherniak, A.; Vazquez, F.; Montgomery, P. G.; Weir, B. A.; Kryukov, G.; Cowley, G. S.; Gill, S.; Harrington, W. F.; Pantel, S.; Krill-Burger, J. M.;, et al. Defining a Cancer Dependency Map. Cell 2017, 170 (3), 564–576 e516. DOI: 10.1016/j.cell.2017.06.010.

(21) Landrum, M. J.; Lee, J. M.; Benson, M.; Brown, G. R.; Chao, C.; Chitipiralla, S.; Gu, B.; Hart, J.; Hoffman, D.; Jang, W.;, et al. ClinVar: improving access to variant interpretations and supporting evidence. Nucleic Acids Res 2018, 46 (D1), D1062–D1067. DOI: 10.1093/nar/gkx1153.

(22) Pedregosa, F. V., Gäel; Gramfort, Alexandre; Michel, Vincent; Thirion, Bertrand; Grisel, Olivier; Blondel, Mathieu; Prettenhofer, Peter; Weiss, Ron; Dubourg, Vincent; Vanderplas, Jake; Passos, Alexandre; Cournapeau, David; Brucher, Matthieu; Perrot, Matthieu; Duchesnay, Édouard. Scikit-learn: Machine Learning in Python. J Mach Learn Res 2011, 12, 2825–2830.

(23) Chen, T.; Guestrin, C. XGBoost. In Proceedings of the 22nd ACM SIGKDD International Conference on Knowledge Discovery and Data Mining, 2016.

(24) Ke, G. M., Qi; Finley, Thomas; Wang, Taifeng; Chen, Wei; Ma, Weidong; Ye, Qiwei; Liu, Tie-yan. LightGBM: A Highly Efficient Gradient Boosting Decision Tree. In Advances in Neural Information Processing Systems 30 (NIPS 2017), 2017.

(25) Guney, E.; Menche, J.; Vidal, M.; Barabasi, A. L. Network-based in silico drug efficacy screening. Nat Commun 2016, 7, 10331. DOI: 10.1038/ncomms10331.

(26) Wang, S.; Sun, H.; Liu, H.; Li, D.; Li, Y.; Hou, T. ADMET Evaluation in Drug Discovery. 16. Predicting hERG Blockers by Combining Multiple Pharmacophores and Machine Learning Approaches. Mol Pharm 2016, 13 (8), 2855–2866. DOI: 10.1021/acs.molpharmaceut.6b00471.

(27) Kuhn, M.; Letunic, I.; Jensen, L. J.; Bork, P. The SIDER database of drugs and side effects. Nucleic Acids Res 2016, 44 (D1), D1075–1079. DOI: 10.1093/nar/gkv1075.

(28) Chen, M.; Suzuki, A.; Thakkar, S.; Yu, K.; Hu, C.; Tong, W. DILIrank: the largest reference drug list ranked by the risk for developing drug-induced liver injury in humans. Drug Discov Today 2016, 21 (4), 648–653. DOI: 10.1016/j.drudis.2016.02.015.

(29) Brown, A. S.; Patel, C. J. A standard database for drug repositioning. Sci Data 2017, 4, 170029. DOI: 10.1038/sdata.2017.29.

(30) Orphanet: an online database of rare diseases and orphan drugs. Orphanet. https://www.orpha.net (accessed June 2026).

(31) Vasilevsky, N. A.; Matentzoglu, N. A.; Toro, S.; Flack, J. E.; Hegde, H.; Unni, D. R.; Alyea, G. F.; Amberger, J. S.; Babb, L.; Balhoff, J. P.;, et al. Mondo: Unifying diseases for the world, by the world. medRxiv 2022. DOI: 10.1101/2022.04.13.22273750.

(32) Subramanian, A.; Narayan, R.; Corsello, S. M.; Peck, D. D.; Natoli, T. E.; Lu, X.; Gould, J.; Davis, J. F.; Tubelli, A. A.; Asiedu, J. K.;, et al. A Next Generation Connectivity Map: L1000 Platform and the First 1,000,000 Profiles. Cell 2017, 171 (6), 1437–1452 e1417. DOI: 10.1016/j.cell.2017.10.049.

(33) Corsello, S. M.; Bittker, J. A.; Liu, Z.; Gould, J.; McCarren, P.; Hirschman, J. E.; Johnston, S. E.; Vrcic, A.; Wong, B.; Khan, M.;, et al. The Drug Repurposing Hub: a next-generation drug library and information resource. Nat Med 2017, 23 (4), 405–408. DOI: 10.1038/nm.4306.

(34) Turei, D.; Korcsmaros, T.; Saez-Rodriguez, J. OmniPath: guidelines and gateway for literature-curated signaling pathway resources. Nat Methods 2016, 13 (12), 966–967. DOI: 10.1038/nmeth.4077.

(35) Minikel, E. V.; Karczewski, K. J.; Martin, H. C.; Cummings, B. B.; Whiffin, N.; Rhodes, D.; Alfoldi, J.; Trembath, R. C.; van Heel, D. A.; Daly, M. J.;, et al. Evaluating drug targets through human loss-of-function genetic variation. Nature 2020, 581 (7809), 459–464. DOI: 10.1038/s41586-020-2267-z.

(36) Feijoo, F.; Palopoli, M.; Bernstein, J.; Siddiqui, S.; Albright, T. E. Key indicators of phase transition for clinical trials through machine learning. Drug Discov Today 2020, 25 (2), 414–421. DOI: 10.1016/j.drudis.2019.12.014.

(37) Huang, K.; Chandak, P.; Wang, Q.; Havaldar, S.; Vaid, A.; Leskovec, J.; Nadkarni, G. N.; Glicksberg, B. S.; Gehlenborg, N.; Zitnik, M. A foundation model for clinician-centered drug repurposing. Nat Med 2024, 30 (12), 3601–3613. DOI: 10.1038/s41591-024-03233-x.

(38) Abraham, W. T.; Lindenfeld, J.; Ponikowski, P.; Agostoni, P.; Butler, J.; Desai, A. S.; Filippatos, G.; Gniot, J.; Fu, M.; Gullestad, L.;, et al. Effect of empagliflozin on exercise ability and symptoms in heart failure patients with reduced and preserved ejection fraction, with and without type 2 diabetes. Eur Heart J 2021, 42 (6), 700–710. DOI: 10.1093/eurheartj/ehaa943.

(39) Packer, M.; Anker, S. D.; Butler, J.; Filippatos, G.; Pocock, S. J.; Carson, P.; Januzzi, J.; Verma, S.; Tsutsui, H.; Brueckmann, M.;, et al. Cardiovascular and Renal Outcomes with Empagliflozin in Heart Failure. N Engl J Med 2020, 383 (15), 1413–1424. DOI: 10.1056/NEJMoa2022190.

(40) Torres, V. E.; Chapman, A. B.; Devuyst, O.; Gansevoort, R. T.; Grantham, J. J.; Higashihara, E.; Perrone, R. D.; Krasa, H. B.; Ouyang, J.; Czerwiec, F. S.;, et al. Tolvaptan in patients with autosomal dominant polycystic kidney disease. N Engl J Med 2012, 367 (25), 2407–2418. DOI: 10.1056/NEJMoa1205511.

(41) Razuvayevskaya, O.; Lopez, I.; Dunham, I.; Ochoa, D. Genetic factors associated with reasons for clinical trial stoppage. Nat Genet 2024, 56 (9), 1862–1867. DOI: 10.1038/s41588-024-01854-z.

(42) Minikel, E. V.; Painter, J. L.; Dong, C. C.; Nelson, M. R. Refining the impact of genetic evidence on clinical success. Nature 2024, 629 (8012), 624–629. DOI: 10.1038/s41586-024-07316-0. Nelson, M. R.; Tipney, H.; Painter, J. L.; Shen, J.; Nicoletti, P.; Shen, Y.; Floratos, A.; Sham, P. C.; Li, M. J.; Wang, J.; et al. The support of human genetic evidence for approved drug indications. Nat Genet 2015, 47 (8), 856–860. DOI: 10.1038/ng.3314.

(43) Gayvert, K. M.; Madhukar, N. S.; Elemento, O. A Data-Driven Approach to Predicting Successes and Failures of Clinical Trials. Cell Chem Biol 2016, 23 (10), 1294–1301. DOI: 10.1016/j.chembiol.2016.07.023.

