## Supplementary Information for "MERIT: Mechanism driven model predicts drug outcomes and nominates indications for failed drugs"

This document contains all supplementary notes, figures and tables (***Notes S1–S9, Figures S1–S3, Tables S1–S20***). All values are computed on the modeling cohort (3,133 trials, 753 compounds) and the released result files.

### SUPPLEMENTARY NOTES

#### Note S1. Full feature-module enumeration

The per-module counts below are indicative, and the released *feature_list_final.csv* gives the exact module-family assignment for every candidate feature.

**Module 1 - Tissue interaction (97 features)**: Predicted binding scores are weighted by tissue-specific expression from the Human Protein Atlas[1]. Thirteen organ systems profiled (heart, brain, liver, kidney, immune, respiratory, endocrine, gastrointestinal, reproductive, musculoskeletal, skin, adipose, other).

**Module 2 - Network-pathway enrichment** (27 features): Drug binding targets mapped onto STRING[2] protein–protein interaction (PPI) networks. Pathway enrichment (false discovery rate, FDR < 0.05) quantified against Open Targets[3] disease targets (top 50 genes per disease). Features include pathway-disease overlap (count and fraction), protein-protein interaction (PPI) network reachability (fraction of disease genes reachable through drug's interaction network), mechanism classification (immune, apoptosis, cell cycle, epigenetic, DNA damage), and off-target pathway activation.

**Module 3 - Binding specificity** (30 features): Distribution statistics of predicted binding scores across the proteome (percentiles, concentration, entropy) and direct-target engagement summaries.

**Module 4 - Safety pharmacology** (45 features): Predicted binding to known toxicity targets (cardiac ion channels including hERG, hepatic enzymes, renal transporters, DNA damage machinery) and essential housekeeping genes. Per-organ essential gene binding burden.

**Module 5 - Pharmacokinetic predictions** (10 features): DruMAP[4]-predicted intrinsic clearance, fraction unbound (human and rat), brain penetration (Kp, brain), volume of distribution, Caco-2 permeability, and CYP substrate classification (1A2, 2C9, 2D6, 3A4).

**Module 6 - Target-disease biology** (30 features): Target-disease mechanism-of-action and biological-association evidence from ChEMBL[5], Open Targets and ClinGen[6], together with STRING network and KEGG[7] pathway co-membership linking the compound's targets to the disease, computed from the target-indication pair alone (***Note S2***).

**Module 7 - Genetics** (16 features): DepMap[8] dependency and Mendelian/ClinVar[9] germline-genetic support for the target-disease relationship.

**Module 8 - Disease complexity & trial context** (15 feature): Disease category flags, mortality, and trial-level context describe the indication and the trial. Disease complexity is the indication's historical failure rate, computed within each CV training fold using Bayesian smoothing toward global rate, with diseases absent from training assigned the global rate.

**Module 9 - Trial-design context** (15 features): Five features from curated reference tables, and ten protocol features. The endpoint-physiology score measures whether the compound's target controls the physiological process of the primary endpoint. Population eligibility flag records whether the trial required the relevant biomarker for eligibility, or was conducted in an indication where that target is the established cause of disease. The endpoint type feature classifies the primary endpoint as surrogate, functional or clinical outcome. The cardiovascular endpoint mechanism-match applies only to trials whose primary endpoint is a major adverse cardiovascular event, recurrent stroke or venous thromboembolism, and measures whether published evidence supports the target for event reduction. The failure precedent flag measures whether a compound of the same target class had already failed Phase III in an earlier trial of the same indication. The protocol features are number of arms, randomization, masking, single group flag, primary endpoint count, secondary endpoint count, primary endpoint timeframe, eligibility criteria count, comparator type and maximum age. Feature selection excludes this module for the efficacy and overall classifiers, but not for the safety classifier. Each feature must pass a leak check where a feature whose availability alone predicts the outcome at an AUROC of 0.58 or above is excluded (***Table S3***).

### Note S2. Target-disease biology and genetics features

The primary target is the mechanism-of-action annotation with a target identity from ChEMBL. All remaining features are computed on the target-disease pair, not on drug identity.

Mechanism-fit module uses four sources. The first is the Open Targets target-disease association, restricted to its six biology evidence channels, namely genetic association, genetic literature, somatic mutation, RNA expression, animal model and affected pathway, with the clinical and drug-disease co-mention channels excluded. The genetic literature channel reports genetic association described in literature. The remaining three sources are STRING interaction network between the compound's targets and the disease gene module, ClinGen germline-genetic support for the target-disease relationship, and KEGG pathway co-membership. A within-disease rank of each is also computed.

Genetic features come from DepMap dependency of disease models on the target, and Mendelian and ClinVar germline genetic support for the target-disease relationship.

Every feature is a public database value reproducible from the released feature matrix. Three further properties follow from the source data. The genetic-association and Mendelian channels are disease-risk evidence assembled independently of trial outcomes. Each channel is available for every compound in the cohort, so a feature's availability is not associated with the outcome. These features are weakly related to the disease base rate. Restoring the excluded clinical and literature channels changes the whole model, including the reported AUROCs.

These channels use a single current snapshot of each database rather than from releases frozen at each temporal cutoff, which limits the temporal analysis.

### Note S3. Benchmark reproduction protocol

#### *S3-1. HINT*

The retrained implementation comprises a message-passing molecular encoder, GRAM embedding of disease ICD codes, BioBERT embedding of eligibility-criteria text and ADMET pretraining, formatted to the required input[10]. Both methods are scored on one shared fold set of 2,787 unique NCTs over 5 seeds × 5 folds. Each NCT is assigned to exactly one-fold per seed, so all rows of a trial testing several indications are assigned together. No trial appears in both the validation split and the training set. Each fold runs 5 epochs after the released ADMET pretraining. The structure-blind evaluation assigns the same 2,787 NCTs to folds at random, using the same fold file HINT's own structure-blind evaluation used.

Two conditions were tested before the comparison was used. The first is whether the implementation is converged rather than under-trained. Retrained from scratch on HINT's own trial outcome prediction (TOP) benchmark dataset, the Phase III split of 1,146 test trials at a 75% base rate, it reached AUROC 0.715 single-pass and 0.730 bootstrap, against the released predictions at 0.721. Ten epochs gave 0.710, so 5 epochs is sufficient for convergence. The second is the effect of the embedding substitution. HINT distributes its eligibility-criteria embeddings as an empty placeholder, so these were regenerated with dmis-lab/biobert-base-cased-v1.1, mean-pooled to 768 dimensions. Scoring the released weights with these substitute vectors gives AUROC 0.678 against 0.721 with the original vectors. This 0.043 difference occurs only when released weights are combined with substitute vectors. Training and testing on the same regenerated embeddings avoid it.

The two cohorts differ in one input. SMILES and eligibility criteria resolve for every trial in both, whereas ICD codes resolve for 57% of this study's Phase III cohort against 100% on original TOP. Restricting to the ICD-complete subset does not change which method scores higher.

*S3-2. TrialBench*

TrialBench's approval label is predicted on the trial-identifier intersection of the two cohorts, with folds grouped by IK14 so no compound appears in both training and test, over 3 seeds × 5 folds[11]. The classifier is a histogram gradient-boosting model (maximum depth 4, 300 iterations, learning rate 0.05) applied identically to every feature set, so the sets differ only in which columns they contain. Five sets are scored, namely MERIT's biology features excluding trial-design and trial-context features, the full TrialBench feature set, TrialBench without the enrollment count, TrialBench without the establishment metadata, and the union of MERIT's biology features with TrialBench minus enrollment. The features excluded from MERIT are module 9 together with the drug count and combination flags. The establishment metadata removed from the TrialBench side is matched on sponsor, agency, oversight, data-monitoring committee, FDA-regulated status, data-sharing, collaborator, responsible-party and funding fields. The main comparison is MERIT's biology features against TrialBench's design features with that metadata removed, and the remaining sets are reported on the original TrialBench terms.

*S3-3. Significance testing*

For the HINT comparison the per-fold difference was tested on the 25 paired AUROCs, which share fold assignments and test sets. A paired t-test and a Wilcoxon rank test were run across the 25 folds. A t-test on the 5 per-seed mean differences was run as the conservative unit, since the 5 seeds reshuffle the same trials and the 25 folds are not independent. All

***Table 1*** benchmark values and the corresponding paired-significance tests were computed by the released benchmark script and reproduced in the accompanying notebook.

*S3-4. Memorization gap configurations*

The gap is measured in two configurations. The full-model configuration ran the full production model on the Phase III cohort under the PASS label. Its 2,942 trial-indication rows come from the same 2,787 trials used in the matched HINT comparison, since some trials test more than one indication. The feature-set configuration ran a single gradient-boosting classifier on the TrialBench intersection of 1,142 trials and 412 compounds under the TrialBench label. In both, the structure-blind evaluation assigns the same trials to folds at random, by StratifiedKFold or by random NCT where MERIT's feature set is used. Every structure-blind fold was verified to place at least one compound in both training and test.

### Note S4. Six-axis PCA stability

Principal component analysis of the molecular feature matrix (578 PASS-cohort drugs, drug-level means) recovers six axes that together account for 59.1% of total variance. Each axis is named for the features most strongly correlated with it. The individual shares are 17.7% for PC1, the number of tissues bound, 16.3% for PC2, predicted binding score, 8.6% for PC3, binding to the single most-bound organ, 7.1% for PC4, mechanism-fit features, 5.3% for PC5, anti-target toxicity, and 4.2% for PC6, which is left unnamed for the reason below. PC1–PC5 reproduce across 100 drug-bootstrap resamples with mean loading correlation $|r|$ of 0.90 to 0.93. PC6 is the least stable at mean $|r| = 0.71$, has the smallest share of variance, and its loadings duplicate those of the PC5 anti-target toxicity axis. PC1–PC5 are treated as named axes.

No principal component corresponds to disease-pathway overlap, so it is reported through three features instead. These are whether the drug has any disease-pathway overlap, the fraction of the drug's enriched pathways that contain disease genes, and the number of disease genes in those pathways. Each feature was compared between outcome groups by Cliff's d, with 95% confidence intervals from 2,000 resamples. All three separate FAIL_SAFETY ($n = 87$) from PASS ($n = 2,558$), at $d = -0.14$ [−0.24, −0.04], −0.20 [−0.33, −0.07] and −0.25 [−0.36, −0.13], and no interval includes zero. The same three features separate FAIL_EFFICACY ($n = 495$) from PASS less strongly, at $d = -0.04$ [−0.08, 0.01], −0.09 [−0.14, −0.03] and −0.05 [−0.11, 0.00], where only the second interval excludes zero.

**Note S5. Prediction confidence gate (prospective registration)**

All 55 novel pairs are registered with a calibrated failure probability. The confidence gate then assigns each pair to one of three groups, confident PASS, confident FAIL, or neither. Only the two confident groups are scored when the trials report, and the third is recorded as unscored. This gives 27 scored predictions (24 PASS, 3 FAIL) and 28 unscored pairs. The two thresholds are computed from the out-of-fold predictions of the canonical model, which is the same model fitted on all cohort rows to score the 55 pairs. Those predictions come from cohort indications, whereas each registered pair is a compound tested in an indication that compound was not tested anywhere in the cohort. The precision measured on the cohort may not apply for these pairs, and the prospective test measures this as well as the predictions themselves.

The two thresholds are set at different precision targets. The PASS threshold reaches 90% precision on held-out data. The FAIL threshold does not reach 90% at any threshold that retains a useful number of pairs, since 90% requires $P\_fail \geq 0.62$ and leaves 20 held-out pairs, so it is set at 85%. A candidate threshold is used only if it selects at least 20 held-out pairs, because a precision of 100% on fewer pairs than this is not informative. The precision achieved is reported for each threshold, not the target.

Two pairs are left unscored for reasons the thresholds do not test. The first is fenofibrate in diabetic retinopathy, a confident FAIL whose indication has few training trials and where neither the efficacy nor the safety sub-model predicts failure. The second is metoprolol in Duchenne muscular dystrophy, excluded before the outcome was known based on other published evidence. It is a confident FAIL assigned on the mismatch between the drug's target and the disease, since ADRB1 is not a dystrophin-pathway driver. Its trial endpoint is change in left ventricular ejection fraction, and beta-blockers preserve ejection fraction in Duchenne cardiomyopathy. The prediction therefore follows the indication's historical failure rate and contradicts the model's own endpoint-physiology feature, which is correct in this case, so the FAIL is likely a false positive. The pair stays in the registered set but is not scored. A confidence score and group are recorded for every prediction.

### Note S6. A published transition model under structure-blind evaluation

inClinico is a multimodal ensemble (omics, indication text, clinical-trial design and small-molecule properties, plus a "target choice" knowledge-graph score) trained on tens of thousands of ClinicalTrials.gov programs[12]. Its validation trains on trials reading out through 2017 and tests on 2018–2021 readouts, so a target's clinical history before 2017 is available when its later trials are scored. Its own subgroup analysis quantifies what that history contributes. On trials whose target has no prior clinical use, AUROC decreases from 0.882 to 0.724 and the target-choice score from 0.841 to 0.697.

The reconstruction and its results are in ***Figure S3***. Its structure-holdout evaluation trains on trials starting up to 2015, tests on 2016–2021, and removes the test compound from training. The count of the pair's Phase 2 trials is retained deliberately, despite including trials that started after the prediction date, so that the reconstruction is scored in the same way as inClinico's own feature set. All other precedent features are computed as of each prediction date and exclude the pair's own trial. The reconstruction includes only Open Targets association and a gene-to-disease module, because MERIT's target-disease biology features are not available at this cohort scale. It therefore identifies which of inClinico's inputs produce its reported value, and does not measure what pre-trial biology can predict.

### Note S7. Disjunctive structure of safety risk

The per-detector out-of-fold scores for the safety-cohort failures were used to test whether the six per-mechanism feature groups, while retaining the six detectors are statistically independent of each other, which is the condition under which safety failure is partly disjunctive.

Among the 85 safety failures, the six detector scores are highly correlated with each other before adjustment (mean absolute pairwise Spearman $\rho = 0.73$). This correlation is attributable to the disease-complexity feature common to all six detectors. When that shared disease and context component are removed from each detector, the axis-specific residuals are independent (mean $|\rho| = 0.38$). The hepatic axis has the weakest correlations with the other five ($\rho$ 0.15–0.41). The highest correlations are between the off-target and organ-distribution detectors (promiscuity-cardiac $\rho$ 0.65, network-tissue 0.51). At a top-5% threshold a detected failure exceeds the threshold on approximately 1.4 detectors on average, and the hepatic axis is the only detector above thresholds for some failures. No single detector is sufficient, hence the six are retained as a decomposition alongside the joint classifier. At 85 positives the pairwise residual correlations have wide sampling error, so the values indicate the general correlation rather than precise effect.

### Note S8. SIDER organ-risk concordance

The toxicity-binding features were compared against SIDER[13] to test whether these rank organ-system risk in the same order as post-market adverse-event distributions. The comparison used 878 profiled drugs by name, and was matched to SIDER independently by name. Of these, 480 (54.7%) matched SIDER. The unmatched drugs are post-2014 approvals, including all PARP inhibitors. The toxicity-binding features were grouped into nine organ-specific panels (cardiac, hepatic, renal, dermatologic, gastrointestinal, musculoskeletal, endocrine, neurological and respiratory), each scoring a drug's predicted risk for that organ system. The per-drug Spearman ρ between predicted organ-z rank and SIDER adverse-event-fraction rank is positive for 43.6% of drugs. The median ρ is -0.04 and the mean top-3 organ overlap is 0.77, against the 0.9 overlap expected by chance for this many organ systems. Organ-level concordance is at chance across all drugs. It differs by drug class and is highest in the classes where adverse events cluster in its target organs. SSRI and SNRI achieve ρ +0.34 (top-3 overlap 1.2), β-blockers +0.19 (1.5), NSAIDs +0.29 (1.3) and tyrosine kinase inhibitors +0.09 (1.2). Each exceeds the 0.9 chance overlap. Statin concordance is negative at ρ = -0.25. The HMG-CoA-pathway features resulted in highest predicted z-ranks for hepatic and endocrine organs, while the highest ranked SIDER organs for statins are musculoskeletal, dermatologic and gastrointestinal. Predicted binding to the musculoskeletal panel separates statins from other drugs at the class-mean level (Cohen d = +1.00 against non-statins). For statins, the metabolic-axis features have higher z-ranks than the musculoskeletal panel.

### Note S9. The confident misclassifications are not recoverable from pre-trial features

Four additional outcome-blind pre-trial feature sets were built and evaluated to test whether the confident misclassifications are missing a pre-trial feature or require information that pre-trial data does not contain. Two sets address the safety misclassifications and two address the efficacy misclassifications. Two criteria were applied to each set. The first is whether it separates the confident misclassifications from correct predictions inside the high-confidence region. The second is how many individual cases change outcome in a full structure-holdoutout retrain. Only the two safety sets were retrained, against a safety classifier at fold-averaged AUROC 0.734 that precedes the configuration reported in the main text. All four tests and their metrics are in ***Table S3***.

*S9-1. Safety residual*

First test mapped each compound's mechanism-of-action targets to the organ-toxicity classes implied by that target's physiology, such as cardiac, hepatic and renal. The mapping uses target biology alone without using any drug identity or trial outcome. Mechanism targets were resolved for 89% of compounds by a preceding target-identity pass. When this feature set was added to the noisy-OR safety classifier, no confident safety misclassification changed outcome and the number of false positives increased. The mapping assigns the on-target organ correctly named, for example cinacalcet to metabolic and endocrine organs. Most compounds implicating an organ complete their trials through managed risk, so the mapping duplicates the disease and drug-class features already in the model.

Second test used reactive-metabolite (haptenation) liability. Established bioactivation structural alerts, defined as SMARTS substructure matches, match the idiosyncratic-hepatotoxicity misclassifications correctly. These are thiophene and methylenedioxyphenyl in sitaxentan, thiophene in duloxetine, and acyl-glucuronide-forming carboxylic acid in fasiglifam. The alerts do not separate failures from passes, since the same reactive groups occur in 130 to 412 compounds that mostly pass, and no confident safety misclassifications changed outcome. A structural alert identifies liability, but whether injury occurs depends on individual patient's HLA allele and immune responses not recorded in trial-level data. This residual requires patient-level information.

*S9-2. Efficacy residual*

Among the 163 confident efficacy misclassifications, 23% have a passing trial elsewhere in the cohort testing the same compound in the same indication. Their molecular and indication features are identical, so only within-trial context could separate them. Two tests were conducted (***Table S3***). First test used a more detailed indication representation, a sentence-embedding of the indication string reduced to 24 principal components. Second test used within-trial design variables for the 63 discordant drug-indication pairs, namely regimen (monotherapy/add-on/combination) and population selectivity, both extracted from the trial protocol without reference to the outcome. The only design variables that separate the misclassifications are comparator type and endpoint type. Both are excluded because they are confounded with how an efficacy failure is detected. A single-arm trial cannot register an efficacy failure, and placebo-controlled trials with posted statistics are detected preferentially. If single-arm trials accumulated in the PASS class they would make that label easier to predict. They do not, since their efficacy-failure rate 0.149 is close to the 0.163 for multi-arm trials. The unpredicted quantity in the efficacy task is the trial-specific effect size, which pre-trial features do not contain.

*S9-3. A prospective direction for the idiosyncratic-hepatotoxicity residual*

This residual depends on patient HLA and not the compound alone. It is addressed by selecting patients before a trial begins, not by classifying compounds after the trials have completed. Three inputs are combined for each compound. These are its haptenation liability from the reactive-metabolite structural alerts, any curated drug–HLA associations[14], and the frequency of the relevant risk allele by ancestry (Allele Frequency Net Database[15]). For a compound with a curated association, the risk allele and the ancestries in which it is commonly found in are extrapolated. Four such compounds and associated alleles are abacavir with HLA-B*57:01, carbamazepine with HLA-B*15:02, allopurinol with HLA-B*58:01, and lapatinib with HLA-DRB1*07:01[14, 16]. For a haptenating compound with no curated association, the two common class-II alleles HLA-DRB1*07:01 and *15:01 are nominated as screening candidates, with their ancestry frequencies. No HLA association is published for fasiglifam or sitaxentan, so this is a screening hypothesis and not a validated prediction. It is suggested as a trial-design criteria and not part of the retrospective model.

*S9-4. The safety features, not the number of safety failures, set limit*

Low safety AUROC can be attributed to the small number of safety failures, or that the molecular features do not represent the mechanisms by which these drugs fail. To distinguish these, the safety-failure set was expanded with additional drug-attributable trial terminations of cohort compounds, retrieved from ClinicalTrials.gov and blind-audited under the same protocol, with toxicity attributed explicitly to other combination drugs where applicable. The prior keyword categorization of these candidates was 81% over-inclusive. Most candidates were either toxicity attributable to a combination drug, or terminations for business or enrollment reasons. The audit retained 32 drug-attributable failures. The fold-averaged safety AUROC decreased from 0.734 to 0.709 with unchanged fold variance. The model assigned the added failures lower predicted failure probabilities than the original set, at a mean of 0.43 against 0.51. These occurred through renal, central-nervous-system, vascular and idiosyncratic hepatic mechanisms. Adding labelled failures of these mechanisms did not improve the safety AUROC. This does not remove the power limitation. At 117 positives, the task has approximately 6 events per variable, so the expansion shows that further failures of the mechanisms it recovered do not improve the safety AUROC. A larger safety-failure set spanning more mechanisms might still do so.

*S9-5. Mechanism of the confident safety misclassifications*

A blind per-case audit assigned a clinical mechanism to every confident safety misclassification, using ClinicalTrials.gov together with the drug label and the literature. Most are on-target pharmacology or established class effects. Examples include PPARγ/Nrf2 fluid retention (bardoxolone), multichannel-antiarrhythmic proarrhythmia (dronedarone), AKT-pathway hyperglycemia (an AKT inhibitor), vascular-disrupting cardiovascular toxicity (fosbretabulin), kinase-inhibitor arterio-occlusion (nilotinib) and immunomodulatory-imide second malignancy (lenalidomide). Each shares its molecular target with cohort drugs that passed their trials, for example pioglitazone for PPARγ, amiodarone and 33 other agents for the antiarrhythmic channels, other AKT inhibitors, and imatinib for ABL1. A feature computed from the target or the drug class therefore assigns the failure and its safe counterpart identical values. A second group is attribution error, comprising a transplant-conditioning regimen and a chemoradiotherapy backbone in which the labelled drug is not the cause. These are excluded from the failure set. For a minority of the misclassifications the cause is not a compound property, namely sitaxentan's fatal idiosyncratic hepatotoxicity and first-in-class compounds with no prior basis for prediction.

Across the 607 cohort compounds matched to ChEMBL (95.2%), the predicted safety-failure score separated in-trial safety failures (n = 73) at AUROC 0.801, but separated neither black-box warning (n = 216) nor post-market withdrawal (n = 19). A drug with a black-box warning may still complete its trials under managed risk.

*S9-6. Safety decision layer*

This decision layer is applied after prediction, not inside the classifier. Each indication is assigned one of three risk-tolerance tiers (life-threatening, serious-chronic, symptomatic/benign), scored blind to outcome. A tier-specific operating threshold is then applied to the retained per-axis detector layer (***Table S8***). Lowering the thresholds for the two lower-risk tiers changes the operating point only, and the AUROC is unchanged. False positives among passing trials decrease from 22% to 13% at near-constant sensitivity (recall 42% to 38%) and the remaining false positives are concentrated in managed-risk oncology.

A separate class-based rule corrects a systematic over-prediction. Taxanes account for 17 of the 44 confident fail-safety predictions and all 17 pass. The most promiscuous molecules with the highest toxicity-binding are ranked highest by a structure-based model, thus these compounds are over-predicted. The rule sets an upper limit of 0.15 on the predicted failure probability for cytotoxic drugs (ATC L01A-D), using the outcome-blind ATC class flag. The base-rate supports this limit, since cytotoxic drugss fail for safety in 2.6% of trials against 3.2% cohort-wide. This removes 19 of the 34 confident false positives and increases confident-region precision 0.23 to 0.40, with no loss of confident true failures and no change to the reported AUROC. It is the safety counterpart of the equivalent rule for oncology cytotoxic monotherapy in the efficacy task.

*S9-7. Drug-induced transcriptional response adds no signal*

Neither connectivity reversal nor mechanistic reconstruction separated matched from mismatched drug-disease pairs at any seed threshold or restart rate on either STRING or the OmniPath network. The number of network connections of a compound's targets alone matched or exceeded the AUROC obtained after propagation, which indicates that the mechanistic reconstruction's score reflects network topology rather than the transcriptional measurement.

### SUPPLEMENTARY FIGURES

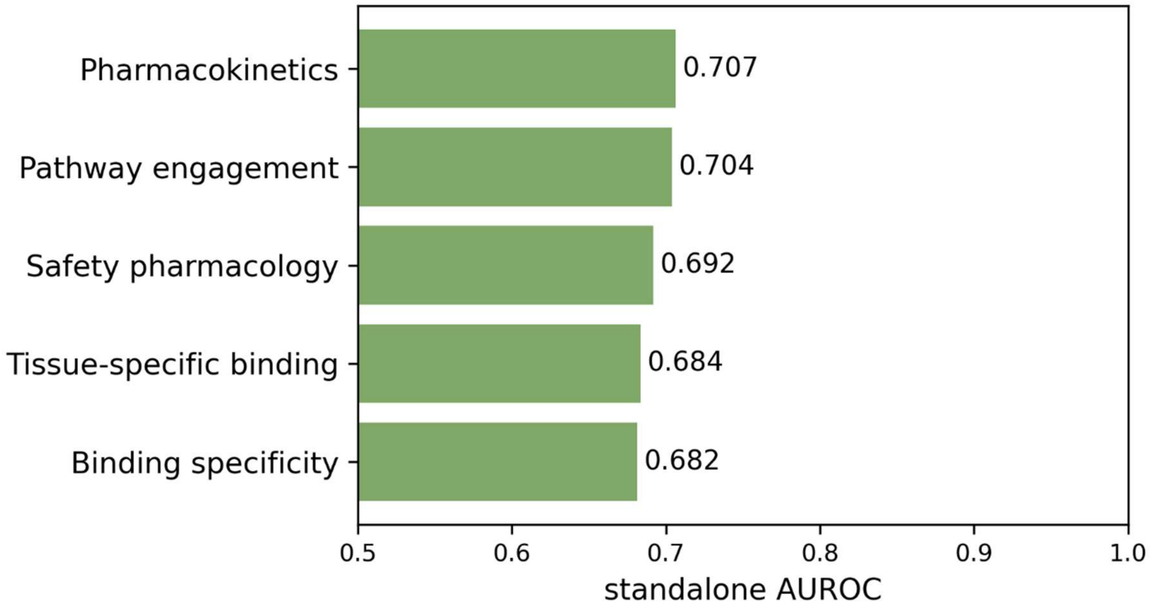


**Figure S1.** No single feature module reaches the full model's predictive accuracy on its own. Standalone overall AUROC (PASS versus any failure) for each of five compound-level feature modules trained on its own, under the same structure-based structure-holdout cross-validation as the full model. Each module scores well below the combined model, so the predictive accuracy comes from combining them rather than any single module.

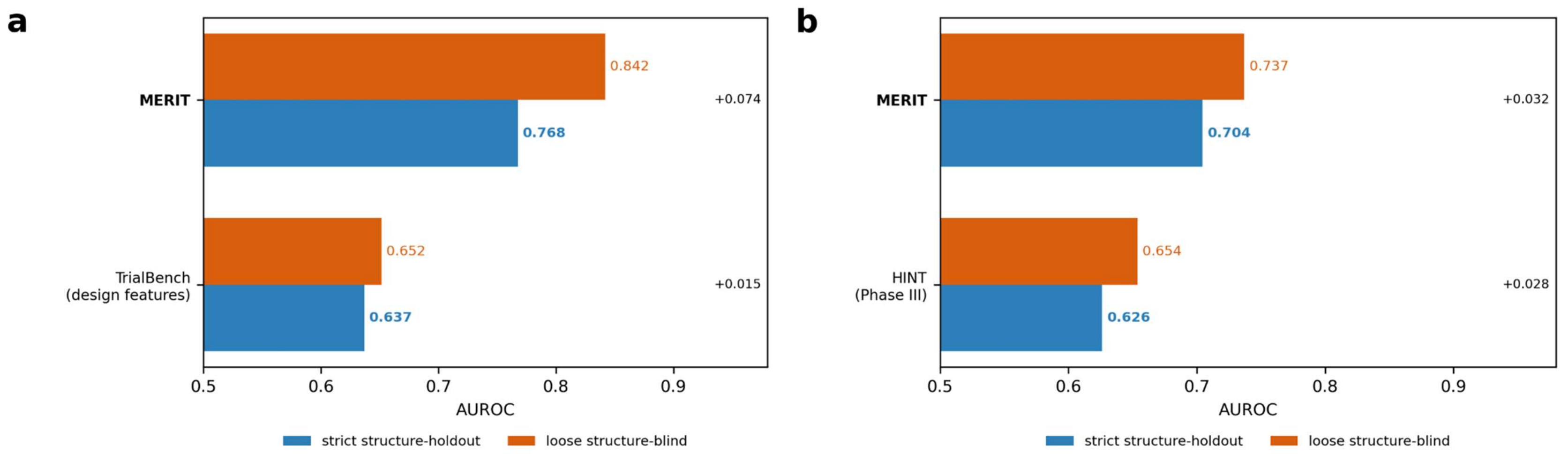


**Figure S2.** Structure-blind evaluation returns a higher AUROC than structure-holdout evaluation. The same models and feature sets were evaluated two ways that differ only in fold assignment. Structure-holdout folds are grouped by InChIKey connectivity layer, so no compound appears in both training and test (blue). Structure-blind folds assign the same trials at random, so a compound's other trials may remain in training (orange). Every method returns a higher AUROC under the structure-blind evaluation. The increment is largest for the mechanism-based features, which assigns one fixed profile per drug. **(a)** Drug-approval benchmark (TrialBench): TrialBench's design features, after removing the enrollment count and establishment metadata, increase from 0.637 to 0.652 (+0.015), and MERIT's pre-trial biology features increase from 0.768 to 0.842 (+0.074). **(b)** Phase III trials: MERIT increases from 0.704 to 0.737 (+0.033) and a retrained HINT increases from 0.626 to 0.654 (+0.028).

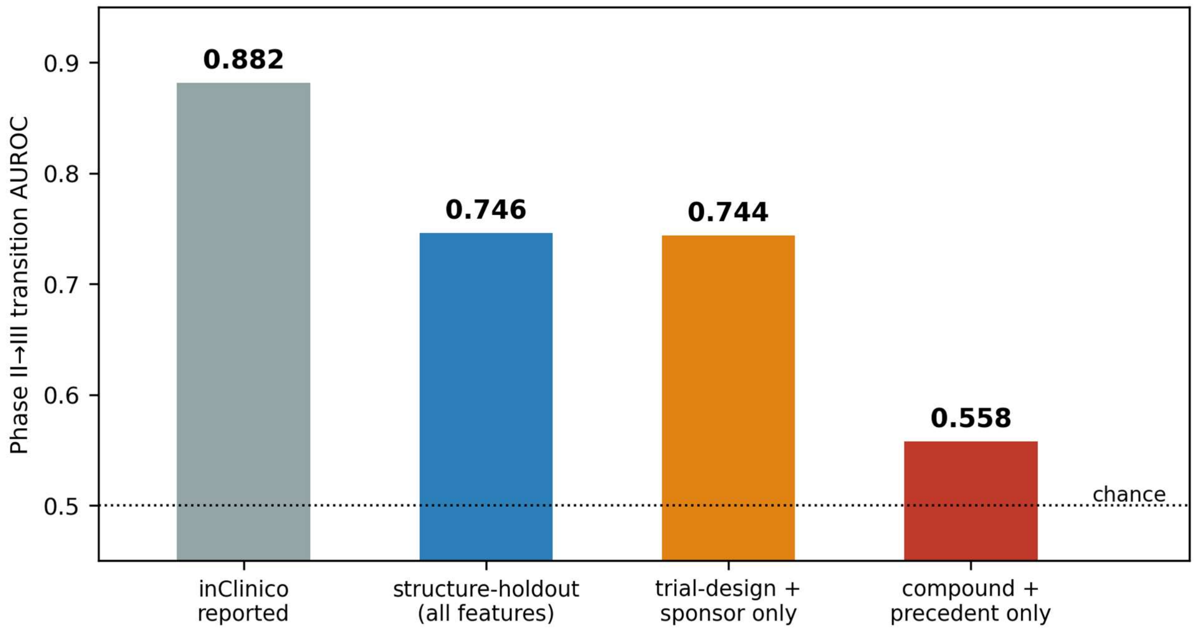


**Figure S3.** In a reconstruction of the Phase II-to-III transition task, predictive accuracy comes from trial-design and sponsor variables rather than from compound properties. inClinico's model and feature set are not public, so a model of the same class was built on 57,043 compound-indication pairs spanning 3,419 small molecules extracted from the AACT registry (***Note S6***). The reported AUROC 0.882 is shown for reference. The reconstruction scores 0.875 under inClinico's compound-grouped random cross-validation, while a structure-holdout evaluation resulted in AUROC of 0.746. The trial-design and sponsor block alone scores 0.744, and its strongest single variable is a count of the pair's Phase 2 trials (univariate AUROC 0.790). This count includes trials that started after the prediction date, and the transition rate increases from 3% at one trial to 48% at four or more, so it is excluded from MERIT's feature set, as are the other reverse-causation variables. With the count and the remaining trial-level variables removed, compound and precedent features (target–disease genetics, mechanism, structural analog and target precedent) predict transition at chance (0.558). This registry-defined transition label differs from the audited efficacy-failure label used in the main text.

### SUPPLEMENTARY TABLES

**Table S1.** Efficacy-evaluation exclusions

| Exclusion category | Trials | Compounds |
|---|---|---|
| Anti-pathogen (target is a pathogen protein) | 321 | 73 |
| Procedural / anesthetic | 95 | 36 |
| Endogenous ligand (binding is normal physiology) | 81 | 13 |
| Mispaired supportive care | 18 | 13 |
| Healthy-volunteer (no disease target) | 3 | 3 |
| Multi-drug attribution | 3 | 3 |

Categories overlap resulting in 512 trials across 130 compounds.

**Table S2.** Arm-level robustness analysis

| Cohort / task | Trials / arms | Compounds | Overall AUROC | Safety AUROC | Efficacy AUROC |
|---|---|---|---|---|---|
| **Trial level (primary)** | 3,133 trials | 753 | **0.770** | **0.784** | **0.765** |
| Arm level (robustness) | 3,326 arms | 891 | 0.710 | 0.688 | 0.720 |

**Table S3.** Outcome-blind tests of the confident misclassifications

| Feature set tested | Task | Within-region AUROC | Misclassifications flipped on retrain | Effect on AUROC | Residual implied |
|---|---|---|---|---|---|
| Target→adverse-event-class organ-toxicity priors | Safety | — | 0/32 | 0.734 → 0.731 (+9 false alarms) | managed-risk; redundant with disease and drug-class context |
| Reactive-metabolite (haptenation) structural alerts | Safety | 0.516 | 0/32 | 0.734 → 0.732 | HLA-restricted immunogenicity (compound-external) |
| Finer indication representation (text embedding) | Efficacy | 0.552 | N/A | N/A | indication already represented by the disease/trial flags and the disease-complexity feature |
| Within-trial design context (regimen, population) | Efficacy | 0.536 | N/A | N/A | trial-specific effect size (compound-external) |

Within-region AUROC = discrimination inside the high-confidence region (chance 0.50); N/A = not applicable (the efficacy feature sets were not retrained).

**Table S4.** Classification metrics by task and operating point

| Threshold | AUROC | PR-AUC | Accuracy | Balanced accuracy | Precision | Recall | F1 | MCC |
|---|---|---|---|---|---|---|---|---|
| *Overall* | | | | | | | | |
| 0.50 | 0.761 ± 0.026 | 0.434 ± 0.044 | 0.829 ± 0.021 | 0.554 ± 0.025 | 0.729 ± 0.191 | 0.118 ± 0.055 | 0.197 ± 0.083 | 0.238 ± 0.076 |
| base rate | 0.761 ± 0.026 | 0.434 ± 0.044 | 0.673 ± 0.089 | 0.692 ± 0.027 | 0.346 ± 0.080 | 0.717 ± 0.137 | 0.449 ± 0.048 | 0.315 ± 0.048 |
| *Efficacy* | | | | | | | | |
| 0.50 | 0.754 ± 0.027 | 0.422 ± 0.061 | 0.841 ± 0.020 | 0.564 ± 0.026 | 0.763 ± 0.109 | 0.139 ± 0.059 | 0.229 ± 0.080 | 0.273 ± 0.063 |
| base rate | 0.754 ± 0.027 | 0.422 ± 0.061 | 0.636 ± 0.099 | 0.678 ± 0.029 | 0.316 ± 0.103 | 0.739 ± 0.139 | 0.419 ± 0.062 | 0.287 ± 0.058 |
| *Safety* | | | | | | | | |
| 0.50 | 0.784 ± 0.066 | 0.226 ± 0.093 | 0.947 ± 0.022 | 0.583 ± 0.034 | 0.263 ± 0.177 | 0.193 ± 0.064 | 0.203 ± 0.086 | 0.189 ± 0.101 |
| base rate | 0.784 ± 0.066 | 0.226 ± 0.093 | 0.893 ± 0.031 | 0.654 ± 0.084 | 0.144 ± 0.089 | 0.397 ± 0.173 | 0.200 ± 0.100 | 0.188 ± 0.110 |

Mean ± SD over 25 structure-holdoutout evaluations. Base-rate thresholds are 18.4% (overall), 17.5% (efficacy), 3.2% (safety). AUROC and PR-AUC are threshold-independent. MCC = Matthews correlation coefficient; PR-AUC = precision-recall area under curve

**Table S5.** Calibration bins for the overall and efficacy tasks

| P(fail) bin | Trials | Observed fail rate | Mean predicted probability |
|---|---|---|---|
| *Overall* | | | |
| [0.00, 0.05) | 82 | 1.2% | 0.041 |
| [0.05, 0.10) | 850 | 5.2% | 0.075 |
| [0.10, 0.20) | 1045 | 10.0% | 0.151 |
| [0.20, 0.30) | 835 | 29.9% | 0.242 |
| [0.30, 0.50) | 239 | 47.3% | 0.370 |
| [0.50, 1.00] | 84 | 77.4% | 0.596 |
| *Efficacy* | | | |
| [0.00, 0.05) | 105 | 5.5% | 0.037 |
| [0.05, 0.10) | 737 | 5.9% | 0.079 |
| [0.10, 0.20) | 759 | 12.5% | 0.151 |
| [0.20, 0.30) | 745 | 23.2% | 0.240 |
| [0.30, 0.50) | 206 | 38.1% | 0.358 |
| [0.50, 1.00] | 87 | 74.8% | 0.654 |

Isotonic-calibrated out-of-fold probabilities, seed-averaged. Overall n = 3,133; efficacy n = 2,639. Safety is reported as a ranker and has no calibration table.

**Table S6.** Traditional machine-learning baselines

| Model | AUROC | PR-AUC | Balanced accuracy | Precision | Recall | F1 | MCC |
|---|---|---|---|---|---|---|---|
| *Overall* | | | | | | | |
| Gradient boosting | 0.739 ± 0.026 | 0.437 ± 0.055 | 0.650 ± 0.019 | 0.260 ± 0.029 | 0.840 ± 0.047 | 0.396 ± 0.034 | 0.238 ± 0.029 |
| Random forest | 0.767 ± 0.026 | 0.476 ± 0.054 | 0.652 ± 0.021 | 0.258 ± 0.032 | 0.868 ± 0.045 | 0.397 ± 0.039 | 0.243 ± 0.036 |
| Logistic regression | 0.707 ± 0.030 | 0.396 ± 0.060 | 0.657 ± 0.029 | 0.289 ± 0.041 | 0.712 ± 0.063 | 0.409 ± 0.041 | 0.245 ± 0.047 |
| *Efficacy* | | | | | | | |
| Gradient boosting | 0.722 ± 0.037 | 0.410 ± 0.067 | 0.644 ± 0.031 | 0.251 ± 0.043 | 0.808 ± 0.055 | 0.380 ± 0.050 | 0.222 ± 0.045 |
| Random forest | 0.741 ± 0.039 | 0.444 ± 0.064 | 0.643 ± 0.030 | 0.244 ± 0.044 | 0.851 ± 0.043 | 0.377 ± 0.053 | 0.224 ± 0.047 |
| Logistic regression | 0.665 ± 0.037 | 0.360 ± 0.048 | 0.604 ± 0.035 | 0.242 ± 0.044 | 0.637 ± 0.093 | 0.347 ± 0.051 | 0.160 ± 0.054 |
| *Safety* | | | | | | | |
| Gradient boosting | 0.780 ± 0.052 | 0.260 ± 0.097 | 0.645 ± 0.079 | 0.129 ± 0.103 | 0.435 ± 0.222 | 0.166 ± 0.088 | 0.161 ± 0.094 |
| Random forest | 0.802 ± 0.061 | 0.292 ± 0.110 | 0.698 ± 0.052 | 0.075 ± 0.040 | 0.782 ± 0.144 | 0.132 ± 0.058 | 0.156 ± 0.060 |
| Logistic regression | 0.767 ± 0.065 | 0.213 ± 0.096 | 0.691 ± 0.064 | 0.079 ± 0.030 | 0.669 ± 0.145 | 0.139 ± 0.046 | 0.155 ± 0.059 |

Threshold metrics use each task's base rate.

**Table S7.** External validation of the safety mechanism axes

| External source | n (positives) | Metric | Value |
|---|---|---|---|
| *Cardiac axis* | | | |
| hERG blockade | 105 (65) | confounds-only CV-AUROC (promiscuity + lipophilicity) | 0.559 |
| hERG blockade | 105 (65) | + cardiac binding panel, CV-AUROC | **0.615** (+0.056) |
| hERG blockade | 105 (65) | cardiac residualized on confounds, AUROC [95% CI] | **0.616 [0.504–0.727]** |
| SIDER cardiac-disorder AEs | 438 | Spearman ρ (cardiac axis) | **+0.106** ($P$ = 0.027) |
| SIDER hepatic AEs (specificity control) | 438 | Spearman ρ (cardiac axis) | +0.051 ($P$ = 0.29) |
| *Hepatic axis* | | | |
| DILIrank | 213 (142) | hepatic binding panel, AUROC | 0.53 |
| DILIrank | 213 (142) | general-promiscuity proxy, AUROC (comparison) | 0.61 |

**Table S8.** Risk-tolerance-adjusted safety operating points (decision layer)

| Tier and indications | Trials | Safety-fail % | Threshold | Failures caught | False positives |
|---|---|---|---|---|---|
| **A.** Life-threatening (oncology, severe/refractory infection) | 795 | 7.7% | 0.65 | 25/61 | 155 (21%) |
| **B.** Serious chronic (cardiac, autoimmune, metabolic, CNS) | 1,074 | 1.6% | 0.45 → 0.60 | 8/17 → 7/17 | 143 (14%) → 79 (7%) |
| **C.** Symptomatic/benign (pain, allergy, dermatology, functional) | 771 | 0.9% | 0.25 → 0.50 | 3/7 → 0/7 | 252 (33%) → 104 (14%) |

Arrows show the default → tuned operating point; I  A is unchanged. Failures caught = safety failures flagged / safety failures present.

**Table S9.** Mechanism heterogeneity of safety failures

| **Primary organ liability** | **n** |
|---|---|
| Broad / multi-organ or unspecified dose-limiting toxicity | 53 |
| Hepatic | 11 |
| Mortality, unspecified organ | 6 |
| Cardiovascular | 4 |
| Hematologic | 3 |
| Neurologic / CNS | 1 |
| Dermatologic | 1 |
| Malignancy (second primary) | 1 |
| **Total in-trial safety-failure trials (all classified)** | **80** |

The 5 FAIL_BOTH trials are not included.

**Table S10.** Terminated-trial label audit

| **Single-pass audit category** | **n** |
|---|---|
| *Safety audit category* | |
| Genuine in-trial safety failure | 75 |
| Combination-partner attribution | 13 |
| Efficacy or strategic | 10 |
| Pre-clinical / external | 7 |
| Post-marketing temporal | 4 |
| Administrative | 2 |
| Business / funding | 1 |
| *Efficacy audit category* | |
| Genuine efficacy failure | 287 |
| Combination-partner attribution | 12 |
| Enrollment only | 7 |
| Business / strategic | 6 |
| Safety mislabel | 2 |
| COVID-19 / operational | 1 |

**Table S11.** Predicted-fit validation on later indications

| Cutoff | Nominations (PASS/FAIL) | AUROC [95% CI] | Mann–Whitney *P* | Precision@3 (random null) | Within-drug concordance |
|---|---|---|---|---|---|
| 2016 | 16 (8/8) | 0.78 [0.51–0.98] | 0.032 | 1.00 (0.50) | 2/3 |
| 2017 | 21 (12/9) | 0.78 [0.53–0.96] | 0.018 | 1.00 (0.57) | 3/4 |
| 2018 | 14 (8/6) | 0.73 [0.42–1.00] | 0.091 | 0.67 (0.57) | 4/4 |

**Table S12.** Signal decomposition and model performance

| **Model** | **Overall AUROC** | **Safety AUROC** | **Efficacy AUROC** | **Marginal AUROC increase** |
|---|---|---|---|---|
| Permuted labels (50 shuffles) | 0.498 ± 0.034 | — | — | — |
| Indication/trial context only | 0.615 ± 0.043 | 0.667 ± 0.089 | 0.641 ± 0.041 | — |
| + disease complexity | 0.689 ± 0.026 | 0.629 ± 0.082 | 0.666 ± 0.038 | **+0.074** |
| + disease complexity, within-fold encoding | 0.739 ± 0.026 | | 0.722 ± 0.037 | |
| + disease complexity, as-of-date encoding | 0.731 ± 0.031 | | 0.713 ± 0.035 | |
| + Molecular-mechanism profile | 0.742 ± 0.027 | 0.717 ± 0.107 | 0.717 ± 0.043 | **+0.053** (efficacy +0.051) |
| Molecular-mechanism profile alone (no disease) | 0.707 ± 0.032 | 0.688 ± 0.091 | 0.695 ± 0.031 | — |
| **MERIT** | **0.770 ± 0.026** | **0.784 ± 0.065** | **0.765 ± 0.026** | — |
| <u>*Matched Phase III comparison (same trials and folds)*</u> | | | | |
| HINT[7] (drug + disease + eligibility embeddings) | 0.626 ± 0.034 | — | — | — |
| **MERIT, same Phase III cohort** | **0.704 ± 0.040** | — | — | — |

Mean ± SD over 25 evaluations, except the permuted-labels row, which is over 50 shuffles. The two encoding rows are computed under the same configuration as each other.

**Table S13.** Precision at industry-relevant top-k

| Task | Base rate | P@10 | P@25 | P@50 | P@100 | P@250 |
|---|---|---|---|---|---|---|
| Overall | 18.4% | 0.90 | 0.88 | 0.86 | 0.75 | 0.60 |
| Safety | 3.2% | 0.30 | 0.20 | 0.24 | 0.22 | 0.15 |
| Efficacy | 17.5% | 1.00 | 0.88 | 0.86 | 0.77 | 0.60 |

**Table S14.** Temporal (leave-future-out) validation

| Cutoff | Test trials (fails) | Overall | Efficacy | Safety |
|---|---|---|---|---|
| 2016 | 356 (137) | 0.704 | 0.649 | 0.746 (31)* |
| 2017 | 246 (104) | 0.737 | 0.731 | 0.822 (27)* |
| 2018 | 199 (89) | 0.737 | 0.710 | 0.904 (23)* |

*Safety positives in the test set.

**Table S15.** Efficacy discrimination and safety over-flagging by drug class

| Drug class | n | Fail % | Efficacy AUROC | Confident FP | FP rate |
|---|---|---|---|---|---|
| *Efficacy discrimination by class* | | | | | |
| Statin | 33 | 21 | 0.896 | | |
| Angiotensin-receptor blocker | 24 | 13 | 0.952 | | |
| BTK inhibitor | 24 | 17 | 0.850 | | |
| Antimetabolite chemotherapy | 114 | 6 | 0.854 | | |
| NSAID / COX inhibitor | 49 | 4 | 0.883 | | |
| Kinase inhibitor (-nib) | 124 | 33 | 0.830 | | |
| JAK inhibitor | 68 | 13 | 0.831 | | |
| Biguanide (metformin) | 84 | 11 | 0.696 | | |
| Antipsychotic | 54 | 11 | 0.583 | | |
| SGLT2 inhibitor | 70 | 11 | 0.512 | | |
| Proton-pump inhibitor | 63 | 16 | 0.542 | | |
| *Safety over-flagging by class* | | | | | |
| Taxane | 77 | 0 | | 31 | 40% |
| PARP inhibitor | 15 | 0 | | 5 | 33% |
| Antimetabolite chemotherapy | 113 | 5 | | 0 | 0% |
| Biguanide | 75 | 0 | | 0 | 0% |
| Proton-pump inhibitor | 54 | 0 | | 0 | 0% |
| Corticosteroid | 147 | 0 | | 0 | 0% |

Per WHO INN-stem drug class on the canonical out-of-fold predictions. Efficacy AUROC reported only where n ≥ 20 with both outcomes. Confident FP counts passing trials flagged in the high-confidence safety region. FP = false positive.

**Table S16.** Representative recoveries of known repositionings

| Compound | Failed indication (fit) | Approved indication (fit) |
|---|---|---|
| imatinib | loiasis (0.67) | chronic myeloid leukemia (0.99) |
| atorvastatin | bicuspid aortic valve (0.51) | hypercholesterolemia (0.98) |
| allopurinol | hypertension (0.23) | gout (0.99) |
| tamoxifen | Duchenne muscular dystrophy (0.07) | breast cancer (0.82) |
| pioglitazone | Alzheimer's disease (0.05) | type 2 diabetes (0.99) |

fit = predicted-fit rank

**Table S17.** Representative novel nominations

| Compound | Target | Nominated indication | Genetic support* |
|---|---|---|---|
| etavopivat | PKLR | hereditary anemia | 0.98 |
| galeterone | CYP17A1 / AR | congenital adrenal hyperplasia (orphan) | 0.93 |
| tivantinib | MET | renal cell carcinoma | 0.95 |
| rilzabrutinib | BTK | immune-mediated disease | 0.97 |

*Genetic support is the Open Targets target–disease genetic association score.

**Table S18.** Real-world corroboration of repositioning nominations

| Corroboration level | n | % |
|---|---|---|
| Same-drug registered clinical trial in the nominated disease | 61 | 42% |
| Same drug approved / documented off-label | 1 | 1% |
| Same-target (class) drug in trials/approved for the disease | 36 | 25% |
| Clinical / target-class precedent | 98 | 67% |
| Preclinical / mechanistic literature only | 31 | 21% |
| Clinical / target-class precedent & preclinical / mechanistic | 129 | 88% |
| Genetic/target rationale only (no clinical or preclinical) | 12 | 8% |
| None / evidence against | 5 | 3% |

**Table S19.** Case studies of repositioning nominations

| Compound (target) | Failed indication (this cohort) | Nominated indication | Independent corroboration |
|---|---|---|---|
| Galeterone (CYP17A1 / AR) | Prostate cancer, Phase III ARMOR3-SV terminated for futility (2016) | Congenital adrenal hyperplasia (orphan endocrine) | Abiraterone, a same-target CYP17A1 inhibitor, trialed in classic 21-hydroxylase-deficiency CAH (e.g. NCT02574910) |
| Evacetrapib (CETP) | Cardiovascular outcomes, Phase III ACCELERATE stopped for futility | Hyperlipidemia / dyslipidemia | Same-class CETP inhibitor obicetrapib in Phase III for dyslipidemia; evacetrapib itself lowered LDL-C by ~31% |
| Idasanutlin (MDM2–p53) | Acute myeloid leukemia, Phase III MIRROS, no survival benefit | Solid tumors, including sarcoma | Idasanutlin advanced into advanced-solid-tumor trials; MDM2 inhibitors (e.g. RG7112, milademetan) tested in MDM2-amplified liposarcoma |

**Table S20.** Analysis scripts

| **Analysis** | **Script(s)** |
|---|---|
| Overall classifier training and probability calibration | scripts/retrain_calibrated.py |
| Classification metrics | notebook 00 Stage 6f, computed from the released out-of-fold predictions |
| Precision at top-*k* and risk zones | scripts/strengthening/compute_risk_zones_and_pk.py |
| Traditional ML baselines | scripts/baseline_models_comparison.py |
| Benchmark comparison + significance | scripts/benchmark/benchmark_final_v2.py, matched_star_phase3_split_compare_clean.py, trialbench_split_compare.py |
| Network-proximity baseline | scripts/network_proximity_baseline.py |
| Memorization-gap / inflation exhibit | scripts/benchmark/make_inflation_exhibit.py (Figure S2), make_proof_ladder_exhibit.py (Figure S3) |
| Decision layer: safety risk-tolerance tiers and cytotoxic cap | scripts/strengthening/decision_layer_compose.py, safety_overflag_decision.py |
| Signal decomposition | scripts/decompose_v8_noclass.py |
| Public-only feature decomposition | scripts/decompose_v8_public_cleanmort.py |

| Analysis | Script(s) |
| --- | --- |
| Temporal (leave-future-out) validation | scripts/temporal_leave_future_out.py |
| As-of-date disease complexity | scripts/disease_complexity_asof.py (notebook 03 §1b) |
| Second-rater label audit | scripts/benchmark/build_second_rater_kit.py, score_second_rater.py |
| PCA stability and per-feature Cliff's d | scripts/phase1/make_fig3_v8.py |
| Phase-stratified AUROC | phase_stratified_auc.py |
| Repositioning, model-in-reverse | scripts/phase1/repositioning_model_reverse.py |
| Mechanism-fit validation on later indications | scripts/repositioning_timesplit.py |
| Prospective registration: build | scripts/benchmark/prereg_C_build_mech.py, prereg_C_build_genetics.py, prereg_C_pull_missing_modules.py, prereg_C_lock.py, prereg_C_finalize_clean.py, prereg_C_novelty_filter.py, prereg_C_completeness.py, prereg_C_confidence_gate.py |
| Prospective registration: score at readout | prereg_C_score_at_readout.py (artifacts results/benchmark/prereg_C/) |

References


(1) Uhlen, M.; Fagerberg, L.; Hallstrom, B. M.; Lindskog, C.; Oksvold, P.; Mardinoglu, A.; Sivertsson, A.; Kampf, C.; Sjostedt, E.; Asplund, A.; et al. Proteomics. Tissue-based map of the human proteome. *Science* **2015**, *347* (6220), 1260419. DOI: 10.1126/science.1260419.
(2) Szklarczyk, D.; Kirsch, R.; Koutrouli, M.; Nastou, K.; Mehryary, F.; Hachilif, R.; Gable, A. L.; Fang, T.; Doncheva, N. T.; Pyysalo, S.; et al. The STRING database in 2023: protein-protein association networks and functional enrichment analyses for any sequenced genome of interest. *Nucleic Acids Res* **2023**, *51* (D1), D638-D646. DOI: 10.1093/nar/gkac1000.
(3) Ochoa, D.; Hercules, A.; Carmona, M.; Suveges, D.; Baker, J.; Malangone, C.; Lopez, I.; Miranda, A.; Cruz-Castillo, C.; Fumis, L.; et al. The next-generation Open Targets Platform: reimagined, redesigned, rebuilt. *Nucleic Acids Res* **2023**, *51* (D1), D1353-D1359. DOI: 10.1093/nar/gkac1046.
(4) Kawashima, H.; Watanabe, R.; Esaki, T.; Kuroda, M.; Nagao, C.; Natsume-Kitatani, Y.; Ohashi, R.; Komura, H.; Mizuguchi, K. DruMAP: A Novel Drug Metabolism and Pharmacokinetics Analysis Platform. *J Med Chem* **2023**, *66* (14), 9697-9709. DOI: 10.1021/acs.jmedchem.3c00481.
(5) Zdrazil, B.; Felix, E.; Hunter, F.; Manners, E. J.; Blackshaw, J.; Corbett, S.; de Veij, M.; Ioannidis, H.; Lopez, D. M.; Mosquera, J. F.; et al. The ChEMBL Database in 2023: a drug discovery platform spanning multiple bioactivity data types and time periods. *Nucleic Acids Res* **2024**, *52* (D1), D1180-D1192. DOI: 10.1093/nar/gkad1004.
(6) Rehm, H. L. B., Jonathan S.; Brooks, Lisa D.; Bustamante, Carlos D.; Evans, James P.; Landrum, Melissa J.; Ledbetter, David H.; Maglott, Donna R.; Martin, Christa Lese; Nussbaum, Robert L.; Plon, Sharon E.; Ramos, Erin M.; Sherry, Stephen T.; Watson, Michael S. ClinGen — The Clinical Genome Resource. *N Engl J Med* **2015**, *372*, 2235-2242.
(7) Kanehisa, M.; Goto, S. KEGG: kyoto encyclopedia of genes and genomes. *Nucleic Acids Res* **2000**, *28* (1), 27-30. DOI: 10.1093/nar/28.1.27.
(8) Tsherniak, A.; Vazquez, F.; Montgomery, P. G.; Weir, B. A.; Kryukov, G.; Cowley, G. S.; Gill, S.; Harrington, W. F.; Pantel, S.; Krill-Burger, J. M.; et al. Defining a Cancer Dependency Map. *Cell* **2017**, *170* (3), 564-576 e516. DOI: 10.1016/j.cell.2017.06.010.
(9) Landrum, M. J.; Lee, J. M.; Benson, M.; Brown, G. R.; Chao, C.; Chitipiralla, S.; Gu, B.; Hart, J.; Hoffman, D.; Jang, W.; et al. ClinVar: improving access to variant interpretations and supporting evidence. *Nucleic Acids Res* **2018**, *46* (D1), D1062-D1067. DOI: 10.1093/nar/gkx1153.
(10) Fu, T.; Huang, K.; Xiao, C.; Glass, L. M.; Sun, J. HINT: Hierarchical interaction network for clinical-trial-outcome predictions. *Patterns (N Y)* **2022**, *3* (4), 100445. DOI: 10.1016/j.patter.2022.100445.
(11) Chen, J.; Hu, Y.; Cai, M.; Lu, Y.; Wang, Y.; Cao, X.; Lin, M.; Xu, H.; Wu, J.; Cao, X.; et al. TrialBench: Multi-Modal AI-Ready Datasets for Clinical Trial Prediction. *Sci Data* **2025**, *12* (1), 1564. DOI: 10.1038/s41597-025-05680-8.
(12) Aliper, A.; Kudrin, R.; Polykovskiy, D.; Kamya, P.; Tutubalina, E.; Chen, S.; Ren, F.; Zhavoronkov, A. Prediction of Clinical Trials Outcomes Based on Target Choice and Clinical

Trial Design with Multi-Modal Artificial Intelligence. *Clin Pharmacol Ther* **2023**, *114* (5), 972-980. DOI: 10.1002/cpt.3008.
(13) Kuhn, M.; Letunic, I.; Jensen, L. J.; Bork, P. The SIDER database of drugs and side effects. *Nucleic Acids Res* **2016**, *44* (D1), D1075-1079. DOI: 10.1093/nar/gkv1075.
(14) Whirl-Carrillo, M.; Huddart, R.; Gong, L.; Sangkuhl, K.; Thorn, C. F.; Whaley, R.; Klein, T. E. An Evidence-Based Framework for Evaluating Pharmacogenomics Knowledge for Personalized Medicine. *Clin Pharmacol Ther* **2021**, *110* (3), 563-572. DOI: 10.1002/cpt.2350.
(15) Gonzalez-Galarza, F. F.; McCabe, A.; Santos, E.; Jones, J.; Takeshita, L.; Ortega-Rivera, N. D.; Cid-Pavon, G. M. D.; Ramsbottom, K.; Ghattaoraya, G.; Alfirevic, A.; et al. Allele frequency net database (AFND) 2020 update: gold-standard data classification, open access genotype data and new query tools. *Nucleic Acids Res* **2020**, *48* (D1), D783-D788. DOI: 10.1093/nar/gkz1029.
(16) Mallal, S. P., Elizabeth; Carosi, Fiampiero; Molina, Jean-Michel; Workman, Cassy; Tomažič, Janez; Jägel-Guedes, Eva; Rugina, Sorin; Kozyrev, Oleg; Cid, Juan Flores; Hay, Phillip; Nolan, David; Hughes, Sara; Hughes, Arlene; Ryan, Susanna; Fitch, Nicholas; Thorborn, Daren; Benbow, Alastair. HLA-B*5701 Screening for Hypersensitivity to Abacavir. *N Engl J Med* **2008**, *358*, 568-579.
